# ZDHHC5 targets Protocadherin 7 to the cell surface by a palmitoylation-dependent mechanism to promote successful cytokinesis

**DOI:** 10.1101/2020.05.24.111831

**Authors:** Nazlı Ezgi Özkan Küçük, Berfu Nur Yiğit, Beste Senem Değirmenci, Mohammad Haroon Qureshi, Altuğ Kamacıoğlu, Nima Bavili, Alper Kiraz, Nurhan Özlü

## Abstract

Cell division requires dramatic reorganization of the cell cortex that is primarily driven by the actomyosin network. We previously reported that Protocadherin 7 (PCDH7) enriches at the cell surface during mitosis which is required for building up the full mitotic rounding pressure. Here we showed that PCDH7 gets palmitoylated and interacts with the palmitoyltransferase, ZDHHC5. Both PCDH7 and ZDHHC5 co-localize at the mitotic cell surface, and they translocate to the cleavage furrow during cytokinesis. PCDH7’s localization depends on palmitoylation activity of ZDHHC5. Loss of expression of PCDH7 impairs active RhoA and phospho-myosin levels at the cleavage furrow and increases the rate of multinucleated cells. This work uncovers a palmitoylation-dependent translocation mechanism for PCDH7 and attributes a regulatory role to contributing actomyosin activity during cytokinesis.

## INTRODUCTION

Cell division is central to life driving many vital cellular events such as proliferation, propagation, development, and regeneration (Rieder and Khodjakov, 2003). As the cell progresses into mitosis, its morphology and surface undergo dramatic reshaping. Adherent cells transiently round-up during mitosis to fulfill the geometric demand of cell division for accurate chromosome segregation (Lancaster et al., 2013; Stewart et al., 2011b). As chromosomes segregate during anaphase, contractile ring machinery assembles to physically divide cells into two, and the daughter cells spread back to regain their interphase morphology during cytokinesis (Ramkumar and Baum, 2016). Morphological changes during mitosis and cytokinesis are primarily driven by the reorganization of actomyosin cytoskeleton network and adhesive systems (Cramer and Mitchison, 1997; Eggert et al., 2006; Rosenblatt, 2008).

At the onset of mitosis, ECT2 activates Rho A at the plasma membrane which activates its downstream effectors, formins and ROCK (Rho-associated protein kinase) (Ramkumar and Baum, 2016). A member of formins, mDia1 localizes to the cell cortex during mitosis and promotes cortical actin nucleation and polymerization (Bovellan et al., 2014). ROCK is a serine/threonine kinase that activates myosin through phosphorylation of its two main substrates. Phosphorylation of the myosin-binding subunit of myosin phosphatase (MYPT1) by ROCK inactivates myosin phosphatase (Kimura et al., 1996). ROCK can also directly phosphorylate the myosin regulatory light chain at Ser-19 (Amano et al., 1996) which results in activation of myosin ATPase that stimulates actin crosslinking and actomyosin contractility (Narumiya et al., 2009).

As a cell rounds up, myosin II progressively accumulates at the cell cortex and the amount of myosin at the cortex is correlated with the rounding pressure (Ramanathan et al., 2015). RNAi-based depletion of cortical myosin II substantially impairs the rounding pressure of mitotic cells (Toyoda et al., 2017). Thus, myosin II plays a fundamental role in cortical reorganization during mitosis by promoting cortical tension (Taubenberger et al., 2020). During cytokinesis, communication between the midzone and actin cortex through Rho signaling drives the assembly of actomyosin contractile ring machinery in between segregating chromosomes (Eggert et al., 2006; Wadsworth, 2021).

Protocadherins (PCDHs) are the largest subgroup of cell surface proteins in the cadherin superfamily (Morishita and Yagi, 2007; Nollet et al., 2000). Although identified as adhesion molecules, the adhesive roles of PCDHs are context-dependent. PCDHs can form homophilic and heterophilic interactions that regulate cell-cell adhesion and downstream signaling events in embryonic and adult tissues (Bradley et al., 1998; Chen et al., 2007; Kahr et al., 2013; Kim et al., 1998; Kuroda et al., 2002; Tai et al., 2010). In our previous study, we investigated how cell surface proteins change during cell division and compared the cell surface proteome of interphase and mitosis cells. Our proteomic analysis identified PCDH7 as one of the proteins that are enriched at the mitotic cell surface and retraction fibers. Knockdown of PCDH7 using siRNAs caused a decrease in the mitotic rounding pressure albeit not as strong as myosin II (Ozlu et al., 2015).

Here, we aimed to unravel the underlying mechanism of the cell cycle-dependent localization of PCDH7 and its role in cell division. PCDH7 localizes to the cell surface at the onset of mitosis and concentrates at the cleavage furrow during cytokinesis. A palmitoyltransferase, ZDHHC5, was identified as proximal interactor of PCDH7. ZDHHC5 interacts with PCDH7, and it targets PCDH7 to the mitotic cortex and cleavage furrow. Cell cycle dependent localization of PCDH7 depends on palmitoylation and requires ZDHHC5 catalytic activity. PCDH7 depletion reduces active RhoA and myosin II levels at the cleavage furrow and increases multinucleation rate. We propose that spatiotemporal regulation of PCDH7 through palmitoylation by ZDHHC5 promotes successful cytokinesis in mammalian cells.

## RESULTS

### PCDH7 localizes to the cleavage furrow during cytokinesis

Our previous study revealed that PCDH7 gets enriched at the cell surface as the cell progresses into mitosis (Ozlu et al., 2015). To probe the spatiotemporal regulation of PCDH7, we analyzed its subcellular localization throughout the cell cycle. For this, we used PCDH7-GFP-BAC cell line in which, GFP tagged PCDH7 is expressed under its own promoter using bacterial artificial chromosome (BAC) transgenomics in Hela cells (Poser et al., 2008). Immunostaining of PCDH7-GFP-BAC cells showed that PCDH7 localized to the cell-to-cell contacts during interphase (**Figure 1A**-top panel), enriched at the cell surface, and retraction fibers at the onset of mitosis (**Figure 1A**-middle panel) and concentrated at the contractile ring during cytokinesis (**Figure 1A**-bottom panel). We observed a similar cleavage furrow and contractile ring localization of PCDH7 in HeLa S3 cells expressing PCDH7:GFP (**Figure 1B, Video S1**). These results suggest that the localization of PCDH7 is regulated in a cell cycle-dependent manner. To test whether cytokinesis-specific localization is due to cleavage furrow formation but not due to freshly forming cell-to-cell contact regions of two daughter cells, we utilized the monopolar cytokinesis approach (Hu et al., 2008; Karayel et al., 2018; Ozlu et al., 2010). In monopolar cytokinesis, cells are arrested in monopolar mitosis by using Kinesin-5 inhibitor S-trityl-L-cysteine (STC) followed by induction of monopolar cytokinesis with CDK inhibitor Purvalanol A. Chromosomes stay in one pole while plasma membrane forms a bud-like extension where the cleavage furrow is greatly expanded without a cell-to-cell contact region (**Figure S1A**). Cleavage furrow proteins, Myosin II, Anillin, mDia and RhoA localizes to the bud like extension, and it biochemically mimics the cleavage furrow of bipolar cytokinesis cells (Hu et al., 2008; Karayel et al., 2018; Ozlu et al., 2010). PCDH7 was enriched at the plasma membrane of monopolar mitosis cells (**Figure 1C**-top panel) and in monopolar cytokinesis cells it was concentrated at the budding site, which corresponds to the cleavage furrow (**Figure 1C**-bottom panel). These results suggest that cleavage furrow localization of PCDH7 is independent of its cell-to-cell contact localization.

**Figure 1.**
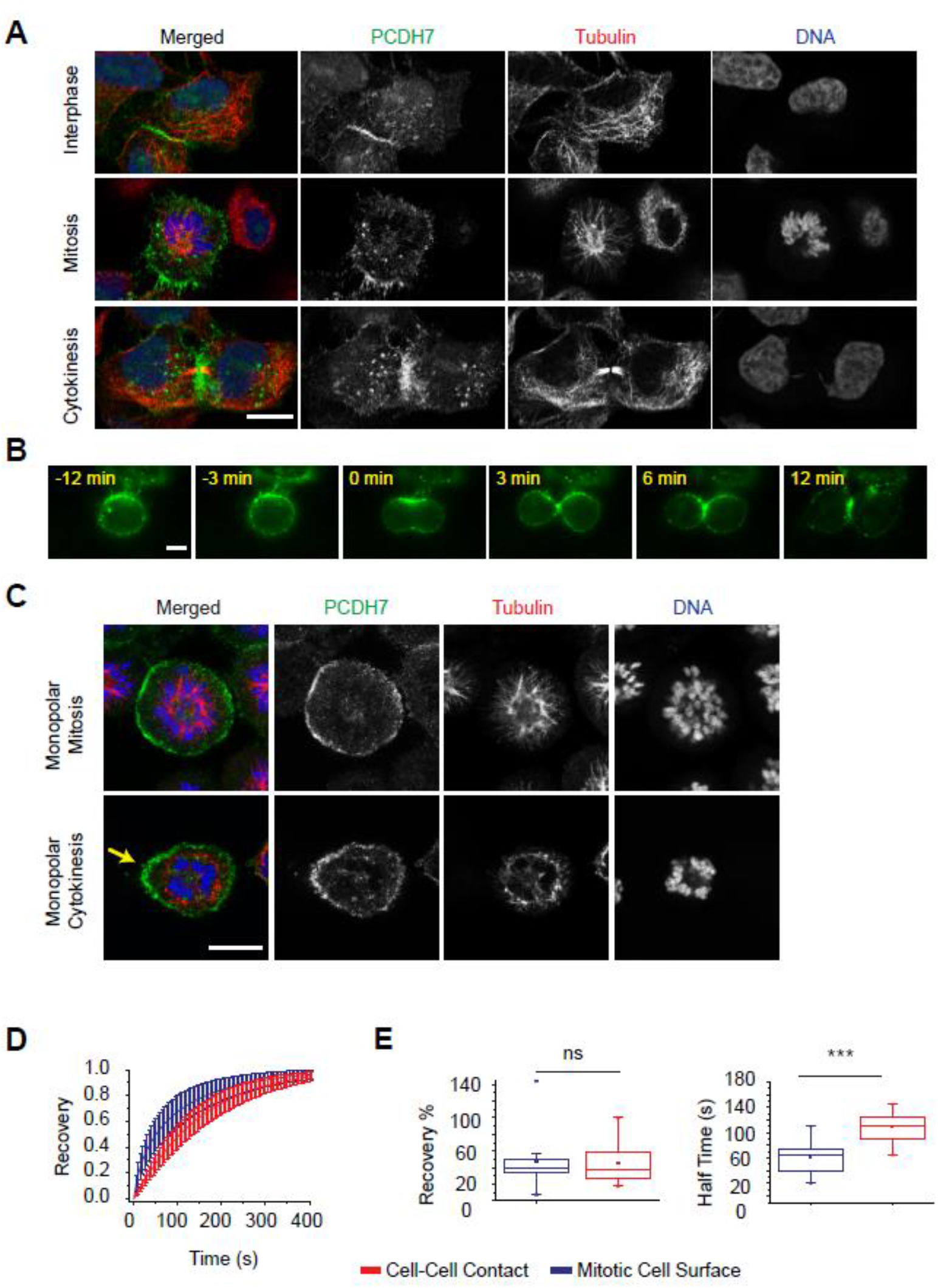
PCDH7 localizes to the mitotic cell surface and cleavage furrow during cell division. **A.** Subcellular localization of PCDH7 (green, anti-GFP), microtubules (red, anti-Tubulin), and DNA (blue, DAPI) in PCDH7-GFP-BAC cells in interphase, mitosis, and cytokinesis. **B**. Live imaging snapshots of PCDH7:GFP expressing HeLa cells during cell division. Relative timing according to cleavage furrow ingression is shown in minutes. **C**. Subcellular localization of PCDH7 (green, anti-GFP), microtubules (red, anti-Tubulin) and DNA (blue, DAPI) in PCDH7-GFP-BAC cells arrested in monopolar mitosis (top panel) and monopolar cytokinesis (lower panel). Yellow arrow indicates the cleavage furrow. **D.** Recovery curve of PCDH7 in mitotic plasma membrane (blue) (n=23) and cell-cell contact regions (red) (n=25) after photobleaching. **E.** Comparison of recovery levels and recovery speeds (halftimes) in mitotic cell surface (blue) (n=23) and cell-cell contact regions (red)(n=25) (right). Statistics used *Mann–Whitney U test*. Scale bars: 10 µm. ***: p<0.001, ns: non-significant

### PCDH7 is more dynamic at the mitotic cell surface than at the cell-cell contact regions

To examine the mitosis dependent translocation of PCDH7, we employed Fluorescence Recovery After Photobleaching (FRAP) experiments. The mobility of PCDH7:GFP in interphase and mitosis cells were compared using PCDH7-GFP-BAC cell line. The GFP signal within the region of interest (ROI) at the plasma membrane of both mitotic (**Figure S1B**-top panel) and interphase (**Figure S1B**-bottom panel) cells were photobleached using a focused laser beam. The fluorescence recovery within ROI was analyzed (**Figure 1D**) using a double normalization algorithm (Phair et al., 2004) and fixed-sized ROI areas (Kappel, 2004). After photobleaching, a significant difference was not observed between the recovery levels of PCDH7:GFP in cell to cell contacts during interphase (average recovery 46.6%) and at the plasma membrane during mitosis (average recovery 45%), as shown in **Figure 1E** (left). However, PCDH7:GFP recovered significantly faster at the mitotic cell surface with an average FRAP half time around 62 seconds, when compared to the cell-to-cell contacts in interphase where the average FRAP half time was around 109 seconds (**Figure 1E**, right). We conclude that PCDH7 is much more dynamic and shows a high turnover rate at the mitotic cell surface in comparison to the cell-to-cell contact regions.

### Depletion of PCDH7 increases the rate of multinucleated cells

Next, we asked about the function of PCDH7 in cell division. Our previous investigation by siRNA-based depletion of PCDH7 revealed that PCDH7 is required for the development of mitotic rounding pressure at the onset of mitosis (Ozlu et al., 2015). To obtain more rigorous data, we utilized the CRISPR-Cas9 genome editing (Ran et al., 2013) to knockout PCDH7 in HeLa cells. PCDH7 was targeted using three individual guide RNAs (sgRNA). Control cells were treated with non-targeting guide RNA (sgNT) in parallel. The knockout of PCDH7 in isolated colonies was validated by Western blot analysis (**Figure S1C**). One colony was selected (PCDH7sg1) to proceed with the phenotypic characterization and non-targeting guide RNA (sgNT) treated cells were used as control. PCDH7 expression levels in PCDH7 knockout (KO), rescue (PCDH7 KO+PCDH7:GFP) and control (PCDH7 KO + eGFP) cells were also validated by Western blotting analysis (**Figure S1D**).

Then, we examined the extent of cell division failures in PCDH7 KO cells. When we analyzed the multinucleation percentage in fixed cells, we observed a moderate but statistically significant increase in the multinucleation rate in PCDH7 KO cells (**Figure 2A-**middle, **Figure 2B-**red bar). The expression of PCDH7:GFP in PCDH7 KO (**Figure 2A**-right) significantly decreased the multinucleation from 3.5 % back to control levels (2%) thus rescued the phenotype **(Figure 2B-**blue bar).

**Figure 2.**
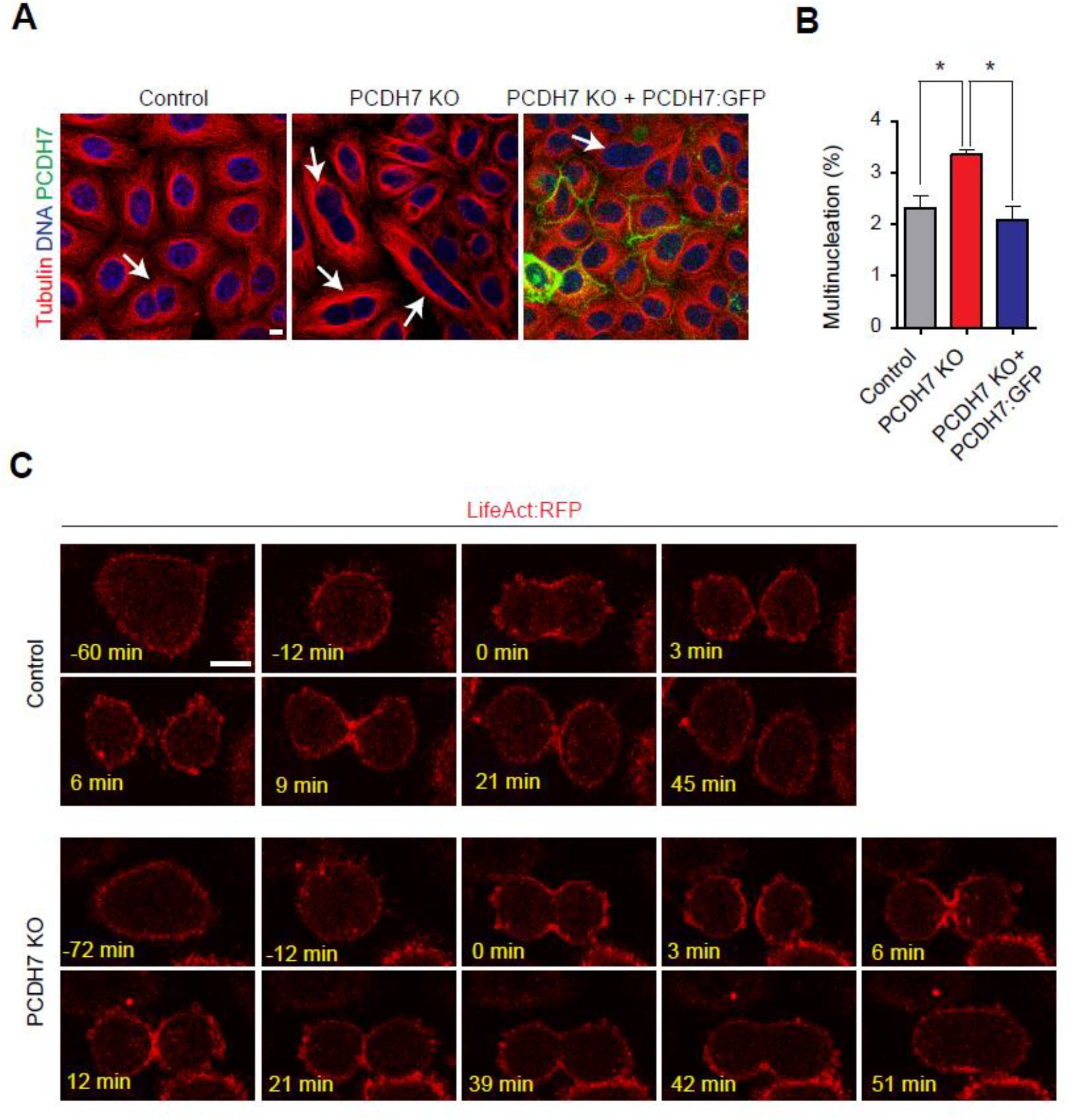
Depletion of PCDH7 increase multinucleation rate and causes cytokinesis failure. **A.** Representative images of the multinucleated cells in control (left) and PCDH7 knock-out (middle), and rescue (PCDH7 KO + PCDH7:eGFP) cells (right). Arrows denote the multinucleated cells. **B.** Quantification of multinucleation percentages of control (2.3%, n=4337, gray), PCDH7 knockout (3.3%, n=4582, red) and rescue cells (PCDH7 KO + PCDH7:eGFP) (2%, n=4591, blue). Statistics used *one-way ANOVA with Tukey’s multiple comparison test*, error bars: SEM **C.** Live imaging snapshots of Control (top panel) and PCDH7 knockout cells (bottom panel) that stably express Actin:RFP. Relative timing according to cleavage furrow ingression is shown in minutes. Scale bars: 10 µm. *: p<0.05

To examine the multinucleation phenotype in PCDH7 KO cells, we performed live cell analysis using cells that stably express Actin:RFP (LifeAct:RFP). In contrast to control cells (**Figure 2C**-top panel, **Video S2**), in PCDH7 KO cells the cleavage furrow ingression started, however, cells were not able to complete cytokinesis and merged back to form multinucleated cells (**Figure 2C**-bottom panel, **Video S3**).

### Actomyosin network and adhesion molecules are among the proximal interaction partners of PCDH7

To address whether the interaction partners of PCDH7 are involved in its cell cycle-dependent translocation, we employed a proteomic approach using the proximity-dependent biotinylation (BioID) method (Roux et al., 2012). In this method, the protein of interest is fused to a promiscuous biotin ligase BirA*, which biotinylates proteins in proximity to the bait, and then the biotinylated proteins are identified using mass spectrometry. Initial immunofluorescence analysis revealed that the PCDH7:BirA* recombinant protein exhibited the expected localization pattern in both mitosis and interphase cells and specifically biotinylated the vicinity **(Figure S2A)**. To biochemically assess the biotinylation efficiency and specificity, we performed streptavidin affinity purification. Western blotting analysis revealed that PCDH7:BirA* exhibited efficient and distinct biotinylation patterns (**Figure S2B**) and streptavidin affinity purification can successfully pull down PCDH7 (**Figure S2C**).

To discover the interphase and mitosis-specific interaction partners of PCDH7, the biotinylated proteins that were isolated from PCDH7 BioID transfected mitotic and interphase cells using streptavidin beads were analyzed in LC-MS/MS. Non-transfected, but biotin supplemented cells were used as a control. The significant interactors were then identified by calculating the spectral count-based fold change between control and PCDH7 BioID cells of 4 biological replicates with a false discovery rate (FDR) of 0.05 (Choi et al., 2015). We identified 78 proteins for mitosis and 129 for interphase cells (**Table S1**), 49 of which were common in both groups. Those proteins were then analyzed using the STRING database (Szklarczyk et al., 2019) and clustered according to GO and KEGG pathway enrichments (Raudvere et al., 2019) using StringApp (Cline et al., 2007; Doncheva et al., 2019). Significant clusters include actomyosin network-related proteins, cell adhesion and cadherin binding proteins, vesicular proteins, and ERM family proteins (**Figure 3A**).

**Figure 3.**
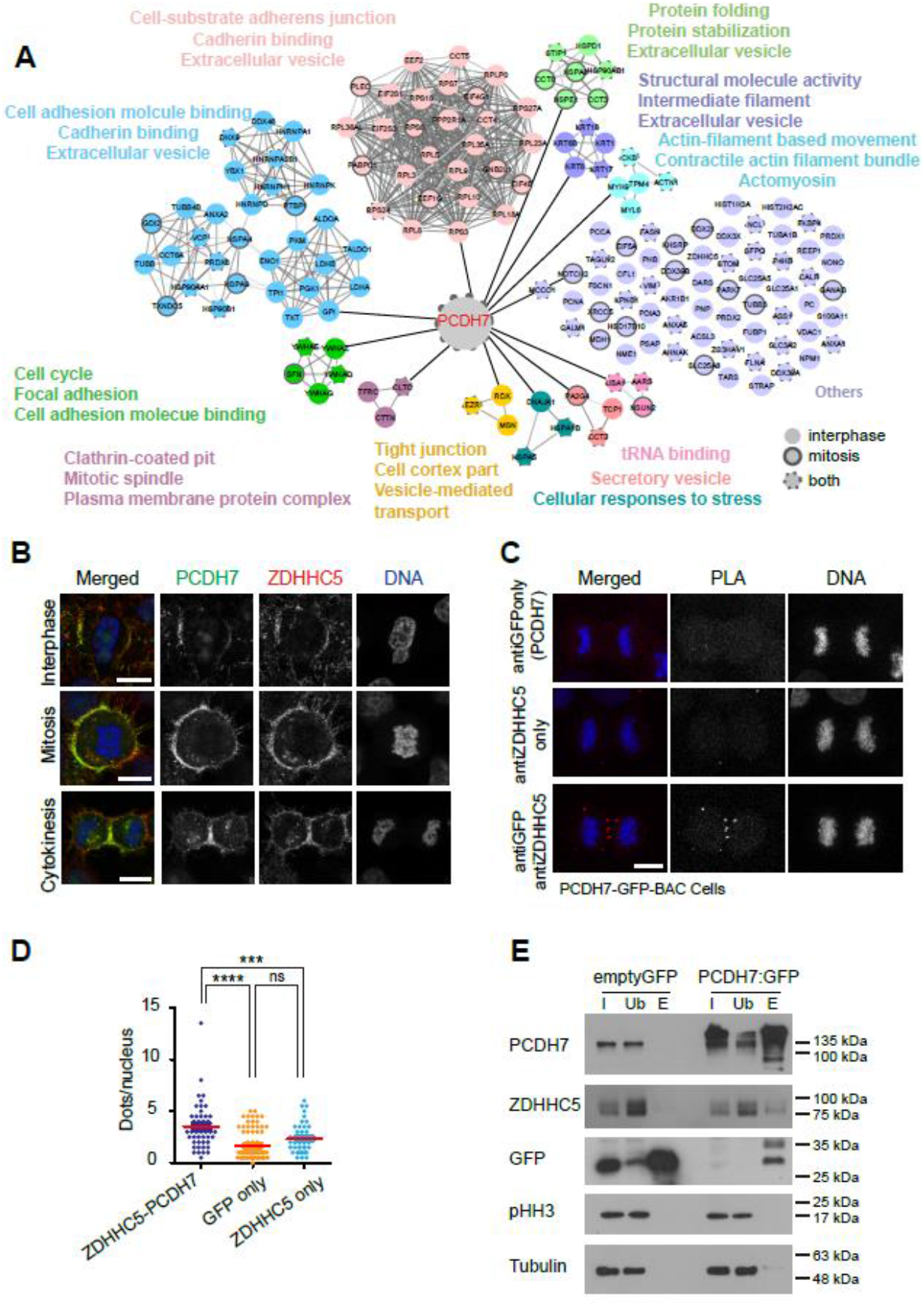
Proximity interactome of PCDH7 in a cell cycle-dependent manner reveals interaction between PCDH7 and ZDHHC5. **A**. Mapping the proximity interaction networking of PCDH7. Different colors represent the significantly enriched clusters after GO and KEGG enrichment analysis. Interphase-specific interactions are represented with non-bordered nodes, mitosis-specific interactors are presented in bordered nodes and common interactors are presented with dashed-bordered nodes. **B**. Subcellular localization of PCDH7 (green, anti-GFP), ZDHHC5 (red, anti-ZDHHC5) and DNA (DAPI, blue) in PCDH7-GFP-BAC cells during interphase (top), mitosis (middle), and cytokinesis (bottom). **C**. Spatial analysis of the interactions between PCDH7 and ZDHHC5 during cytokinesis by in situ proximity ligation assay (PLA). PCDH7-GFP-BAC cells were used for analysis and PCDH7 was targeted using GFP antibody. Control cells were treated with anti-GFP and anti-ZDHHC5 antibody only. The representative images show the interactions between the examined antibody pairs as red fluorescent PLA puncta. DNA is shown in blue (DAPI). **D.** Quantification of the PLA puncta observed in anti-GFP [PCDH7] only antibody-treated cells (n=77), anti-ZDHHC5 only treated cells (n=59) and GFP-ZDHHC5 antibody-treated cells (n=49). Each image is the maximum intensity projection of a Z-stack. The statistic used *one-way ANOVA with Sidak’s multiple comparison test*. **E.** Pull down analysis of PCDH7 using GFP-trap approach. HeLa S3 cells that stably express PCDH7:GFP or empty GFP (control) were arrested in mitosis. Whole-cell lysates (input), unbound fractions (Ub), and elutes (E) were analyzed by Western blotting using PCDH7, GFP, and ZDHHC5 antibodies. Phospho-Histone H3 (pHH3) was used as mitosis marker, α-Tubulin was used as the loading control. I: Input, U: Unbound, E: Elute. Scale bars: 10 µm, ****:p<0.0001, ***: p<0.001, ns: non-significant.

### PCDH7 interacts with Palmitoyltransferase, ZDHHC5, during cell division

Among many interesting candidates, one striking interaction partner of PCDH7 was a palmitoyltransferase (ZDHHC5) that adds the palmityl group to its target proteins. Palmitoylation is a reversible post-translational modification that is known to affect the hydrophobicity, the membrane domain interactions, and the conformation of transmembrane proteins (Blaskovic et al., 2013). To address whether PCDH7 cooperates with ZDHHC5 during cell division, we first visualized subcellular localization of PCDH7 and ZDHH5 in PCDH7-GFP-BAC cells by using anti-GFP and anti-ZDHH5 antibodies respectively. During interphase both PCDH7 and ZDHHC5 localized to the cell-cell contact regions (**Figure 3B**-top panel). During mitosis, both proteins decorated the cell surface and retraction fibers (**Figure 3B**-middle panel) and as cells proceed to cytokinesis, they both accumulated at the cleavage furrow (**Figure 3B**-bottom panel). To further verify their colocalization, we applied the proximity ligation assay (PLA) (Soderberg et al., 2006). For this, we used fixed PCDH7-GFP-BAC cells and anti-GFP and anti-ZDHHC5 antibodies to visualize target proteins’ interaction during cytokinesis (**Figure 3C**). As expected, we observed a significantly higher signal in cells treated with both primary antibodies than the control cells that were treated with only one primary antibody (**Figure 3D**). The PLA signals were accumulated around the cleavage furrow which is in line with the immunofluorescence assay (**Figure 3C**, bottom panel).

Given their co-localization, we tested whether PCDH7 physically interacts with ZDHHC5 during mitosis by performing co-immunoprecipitation using the GFP-trap approach in mitotic PCDH7:GFP expressing cells. We observed that PCDH7 interacts with ZDHHC5 in mitotic cells (**Figure 3E**).

### Palmitoylation of PCDH7 is required for its mitotic cell surface and cleavage furrow localization

Interaction between PCDH7 and ZDHHC5 prompted us to test whether PCDH7 is palmitoylated. To this end, we performed a palmitoylation assay by using Acyl-biotin Exchange chemistry (ABE) (Cline et al., 2007). HeLa S3 PCDH7:GFP expressing cells were used to enrich the PCDH7 protein amount in the samples. Briefly, after performing a high-speed centrifugation-based membrane enrichment, unmodified cysteine thiol groups were blocked using NEM. Half of the sample was treated with HA that cleaves the palmitate groups from cysteines which were then biotinylated using a thiol-reactive biotin molecule. As a negative control, the other half was not treated with HA (HA-). Subsequently, thiol-biotinylated proteins were purified by streptavidin beads (**Figure 4A**). Palmitoylated proteins are expected to be in the hydroxylamine treated (HA+) eluates. Palmitoylation of PCDH7 was tested by Western blotting against anti-PCDH7 antibody. In parallel, antibodies against EGFR and Calnexin were used as positive controls that are known to be palmitoylated. Like EGFR and Calnexin, we observed PCDH7 in the palmitoylated fraction. As expected, PCDH7 was not detectable in the HA-fraction (**Figure 4B**).

**Figure 4.**
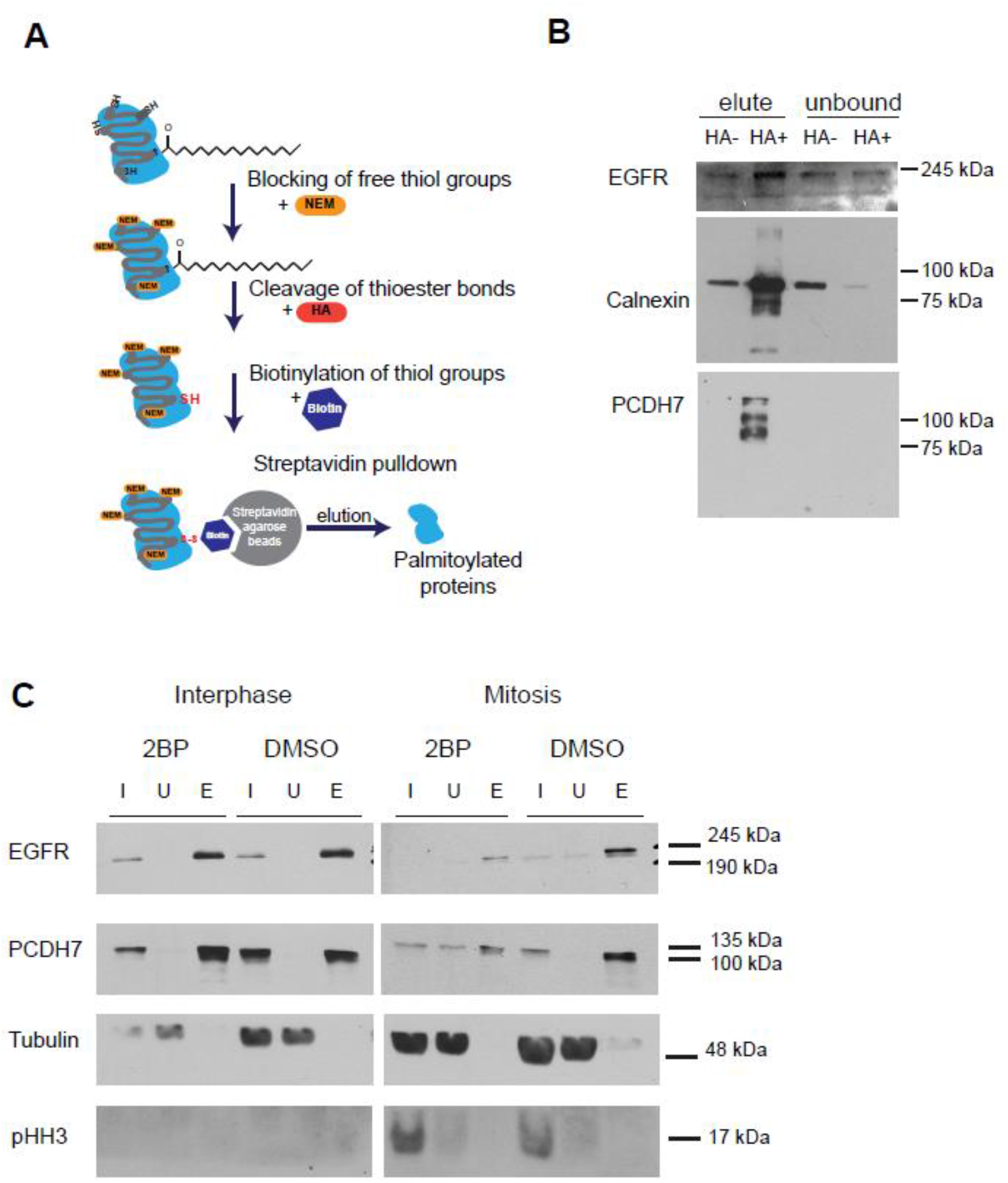
PCDH7 is palmitoylated and palmitoylation is required for cell surface localization of PCDH7. **A.** Schematic illustration of Acyl-biotin Exchange (ABE) chemistry assay to identify palmitoylated proteins. NEM treatment blocks free thiols while Hydroxylamine (HA) treatment cleaves the palmitate groups from cysteine residues those are subsequently biotinylated using a thiol-reactive biotin molecule. Streptavidin pulldown was used to capture the palmitoylated proteins. **B.** Western blot analyses of palmitoylated proteins after ABE assay. HA+ represents hydroxylamine treated sample; HA-represents untreated ones. Proteins that are known to be palmitoylated, EGFR and Calnexin, were used as positive controls. **C.** Western blot analyses of the effect of 2BP inhibitor treatment on cell surface localization of PCDH7. Surface exposed proteins were labeled with a non-permeable amine-reactive biotin and pulled down by streptavidin beads. All fractions (I: Input, U: Unbound, E: Elute) were blotted against EGFR (as a cell surface marker), α-tubulin (as a cytoplasmic marker), and phospho Histone H3 (pHH3) (as a mitotic marker).

To further examine the palmitoylation of PCDH7, we utilized a standard palmitoylation inhibitor, 2-bromopalmitate (2BP) (Webb et al., 2000). We first tested the effect of the palmitoylation inhibitor on PCDH7’s hydrophobicity using TritonX-114 extraction, which separates proteins into detergent and aqueous phases according to their hydrophobic properties (Bordier, 1981). The inhibition of palmitoylation decreased the proportion of PCDH7 in the detergent phase suggesting a reduction in the hydrophobicity of PCDH7 (**Figure S3A**).

To examine the role of palmitoylation in mitosis dependent PCDH7 cell surface localization, we tested the association of PCDH7 to the cell surface after palmitoylation inhibitor (2BP) treatment. For this, we purified cell surface proteins and compared PCDH7 levels in control and 2BP treated cells. Briefly, HeLa S3 cells were treated with 2BP or DMSO (control) overnight, while being synchronized to interphase or mitosis using Thymidine and STC respectively. To enrich for cell surface-exposed proteins, intact cells were labeled with a non-permeable sulfo-NHS-SS-biotin reagent followed by affinity purification of biotinylated proteins using streptavidin (Özkan Küçük et al., 2018). While the 2BP treatment did not affect the biotinylated PCDH7 level in interphase cells (**Figure 4C**-left panel), it strikingly decreased the amount of biotinylated PCDH7 in mitotic cells (**Figure 4C**-right panel). These results suggest that palmitoylation of PCDH7 is required for its mitotic cell surface localization.

To further examine the impact of the palmitoylation inhibitor on the subcellular localization of PCDH7 in mitosis and interphase cells, we performed immunostaining and live imaging of a PCDH7-GFP-BAC cell line treated with 2BP. In line with our previous findings (**Figure 4C**), the inhibition of palmitoylation did not show a noticeable effect on the localization of PCDH7 at the cell-to-cell contacts during interphase **(Figure S3B)**. However, it dramatically affected the surface localization of PCDH7 in mitotic cells (**Figure 5A**) and significantly decreased the plasma membrane enrichment (**Figure S3C**) of PCDH7 (**Figure 5B**). Live imaging of PCDH7 also supported these results (**Figure S3D, Video S4-S5**).

**Figure 5.**
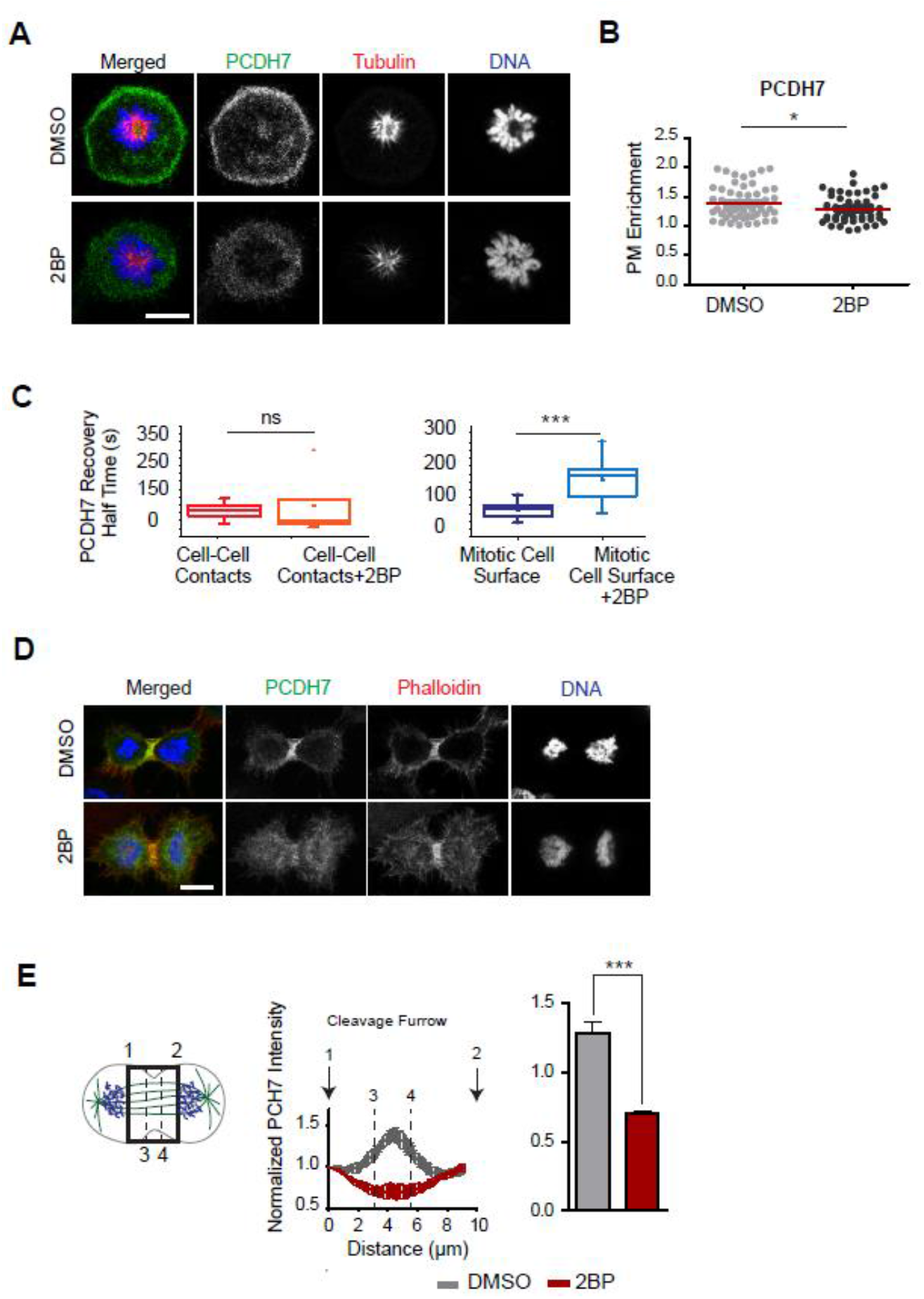
Inhibition of palmitoylation perturbs cell surface and cleavage furrow localization of PCDH7. **A.** PCDH7 (green, anti-GFP) localization at mitosis in control (DMSO) (top) and palmitoylation inhibitor, 2BP treated (bottom) cells those are stably expressing PCDH7-GFP-BAC. Cells are synchronized to monopolar mitosis. Microtubules (anti-β-Tubulin) are shown in red and DNA (DAPI) in blue. **B.** Quantification of the plasma membrane enrichment of PCDH7 in the control (n=56) and 2BP treated (n=54) mitotic cells (right). Statistics used *unpaired two-tailed t-test*. **C.** FRAP analysis of PCDH7 in palmitoylation inhibitor 2BP treated cells. Recovery half times of PCDH7 in cell-cell contacts in 2BP treated (n=6) and control (n=25) cells (left). Recovery half times of PCDH7 in the mitotic cell surface in 2BP treated (n=9) and to control (n=23) cells (right). Statistics used *Mann–Whitney U test*. **D.** PCDH7 (green, anti-GFP) localization at the cleavage furrow in control (top panel) and 2BP treated (bottom panel) PCDH7-GFP-BAC expressing cells. Each image is the maximum intensity projection of a Z-stack. Actin filaments (Phalloidin) are shown in red and DNA (DAPI) in blue. **E.** Illustration of the analyzed region of interests in the cell (left). Intensity profiles (measured from 1 to 2) of PCDH7 in the control (n=13) (gray) and 2BP treated (n=26) (red) cytokinesis cells (middle). The sum of PCDH7 intensities in between the dashed lines covering the +/- 15% distance around the middle zone (measured from 3 to 4), were quantified (right). Intensity profiles were obtained in ImageJ software for the indicated region of interest as previously described (Uretmen Kagiali et al., 2020). Statistics used *unpaired two-tailed t-test*, error bars: SD. Scale bars: 10 µm,*: p<0.05; **: p<0.01; ***: p<0.001 ns: non-significant.

To further investigate the effect of the palmitoylation inhibitor on the dynamic behavior of PCDH7 at the mitotic cell surface, we again took the FRAP approach and measured the extent of PCDH7:GFP recovery at the plasma membrane after photobleaching, in the absence or presence of 2BP. As expected, the inhibition of palmitoylation did not have a significant effect on the FRAP recovery of PCDH7 at the cell-to-cell contacts during interphase (**Figure 5C**-left). On the other hand, 2BP treatment dramatically increased the FRAP recovery time of PCDH7 at the mitotic plasma membrane (**Figure 5C**-right). Finally, we analyzed the effect of 2BP on PCDH7’s localization during cytokinesis (**Figure 5D**). Inhibition of palmitoylation explicitly perturbed the cleavage furrow localization of PCDH7 during cytokinesis (**Figure 5D, E**-left) and led to a decrease in the levels of PCDH7 at the cleavage furrow (**Figure 5E**-right). We conclude that palmitoylation is dispensable for a stable cell-to-cell contact localization of PCDH7 during interphase, but essential for the cortical and cleavage furrow localization of PCDH7 during mitosis and cytokinesis, respectively.

### ZDHHC5 dependent palmitoylation directs PCDH7 to the plasma membrane at the onset of mitosis

Building upon results of palmitoylation-dependent localization of PCDH7 in mitosis and cytokinesis, we next sought to examine the role of ZDHHC5 in this process. To this end, siRNA (**Figure 6A**) or shRNA (**Figure S4A**) mediated knock-down of ZDHHC5 and control (non-targeting shRNA and siRNAs) Hela cells were used. In line with our previous observation, in control siRNA-treated cells, ZDHHC5 and PCDH7 largely co-localize at the mitotic cell surface and at retraction fibers (**Figure 6B**, top panel). Strikingly, ZDHCC5 depletion significantly decreased the PCDH7 signal at the mitotic cell surface (**Figure 6B**, bottom panel, **Figure 6C**). Knockdown of ZDHHC5 using shRNA against ZDHHC5 (**Figure S4A**) also gave similar results (**Figure S4B**). Cell surface localization of PCDH7 during mitosis was significantly decreased upon shZDHHC5 treatment (**Figure S4B**-right).

**Figure 6.**
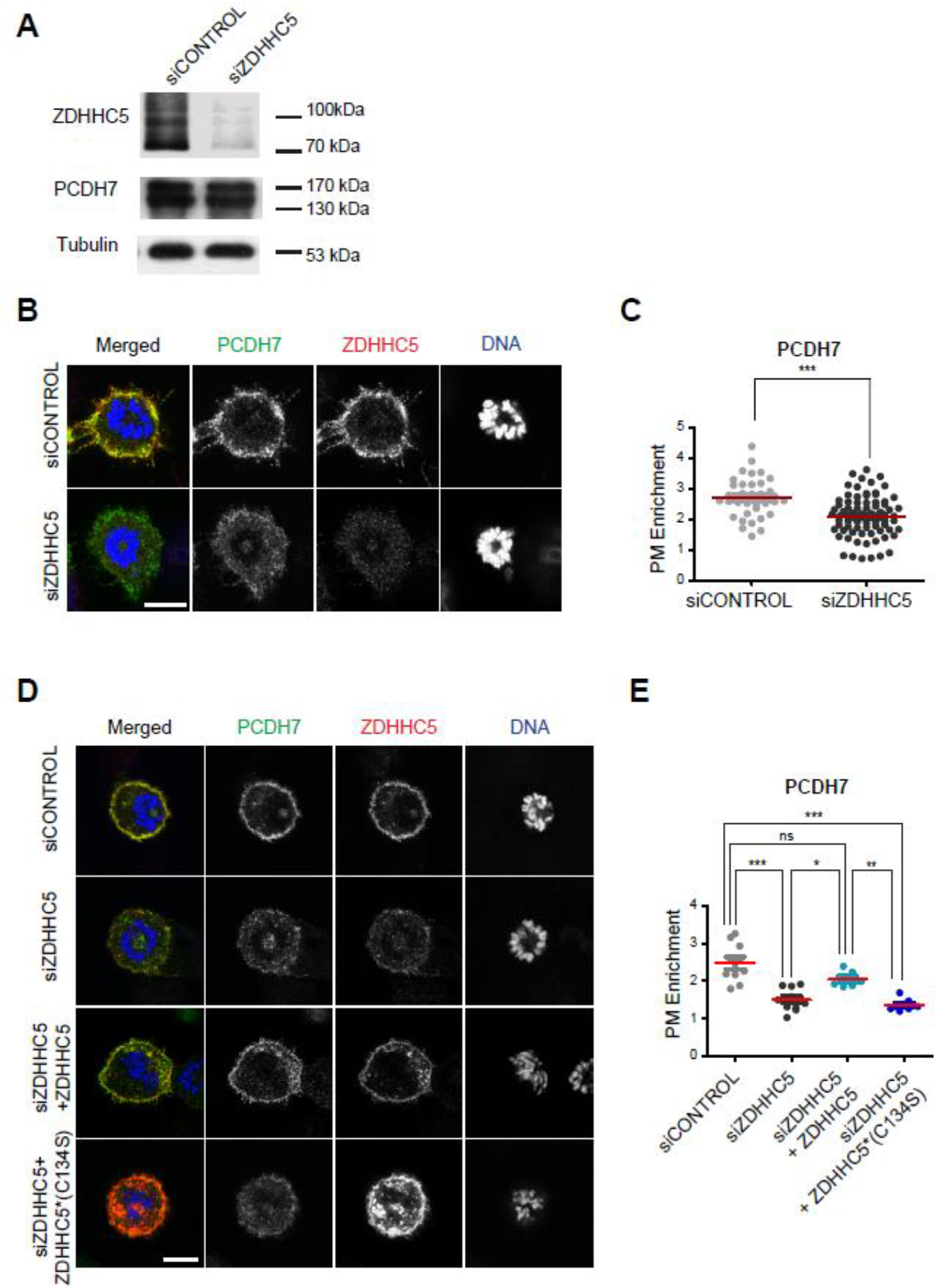
ZDHH5 targets PCDH7 to plasma membrane during mitosis. **A.** Western blot analysis of siZDHHC5 and siCONTROL cells. siZDHHC5 treatment successfully depleted the ZDHHC5 proteins while PCDH7 levels remain unchanged. α-Tubulin was used as the loading control. **B**. PCDH7 (green, anti-GFP) localization in mitotic control (siCONTROL) (top) and ZDHHC5 knockdown (siZDHHC5) (bottom) cells that are stably expressing PCDH7-GFP-BAC. Cells are synchronized to monopolar mitosis. ZDHHC5 is shown in red (anti-ZDHHC5) and DNA in blue (DAPI). **C.** Quantification of the plasma membrane enrichment of PCDH7 during mitosis in the control (n=39) and ZDHHC5 KD (n=83) cells (right). Statistics used *unpaired two-tailed t-test*. **D.** PCDH7 (green, anti-GFP) localization in mitotic control (siCONTROL), ZDHHC5 knockdown (siZDHHC5) and ZDHHC5 rescued (3^rd^ &4^th^ row) cells that are stably expressing PCDH7-GFP-BAC. To restore ZDHHC5 expression, cells were transfected with either wild-type ZDHHC5(3^rd^ row) or catalytically inactive mutant (ZDHHC5*(C134S)) (4^th^ row). **E.** Quantification of the plasma membrane enrichment of PCDH7 during mitosis in the control (n=10) and ZDHHC5 KD (n=10), ZDHHC5 transfected (n=6) and ZDHHC5-mutant (C134S) (n=8) transfected cells (right). Statistics used *one-way ANOVA with Benferonni’s multiple comparison test*. Scale bars: 10 µm, *: p<0.05; **: p<0.01; ***: p<0.001

Next, we tested whether restoring the ZDHHC5 expression can rescue impaired plasma membrane localization of PCDH7 during mitosis. Expression of murine ZDHHC5 in siZDHHC5 cells significantly restored the plasma membrane localization of PCDH7 in mitotic cells. However, catalytically inactive ZDHHC5 (C134S) mutant was not able to rescue this phenotype. PCDH7 levels at the plasma membrane were similar to siZDHHC5 levels (**Figure 6 D, E**). These results indicate that the palmitoylation activity of ZDHHC5 translocates PCDH7 to the plasma membrane at the onset of mitosis.

### ZDHHC5 directs PCDH7 to the cleavage furrow

Next, we examined role of ZDHHC5 in targeting PCDH7 to the cleavage furrow. In ZDHHC5 depleted cells (shZDHHC5), cleavage furrow localization of PCDH7 is perturbed (**Figure 7A**). PCDH7 was no longer enriched at the cleavage furrow in ZDHHC5 knock down cells (**Figure 7B**).

**Figure 7.**
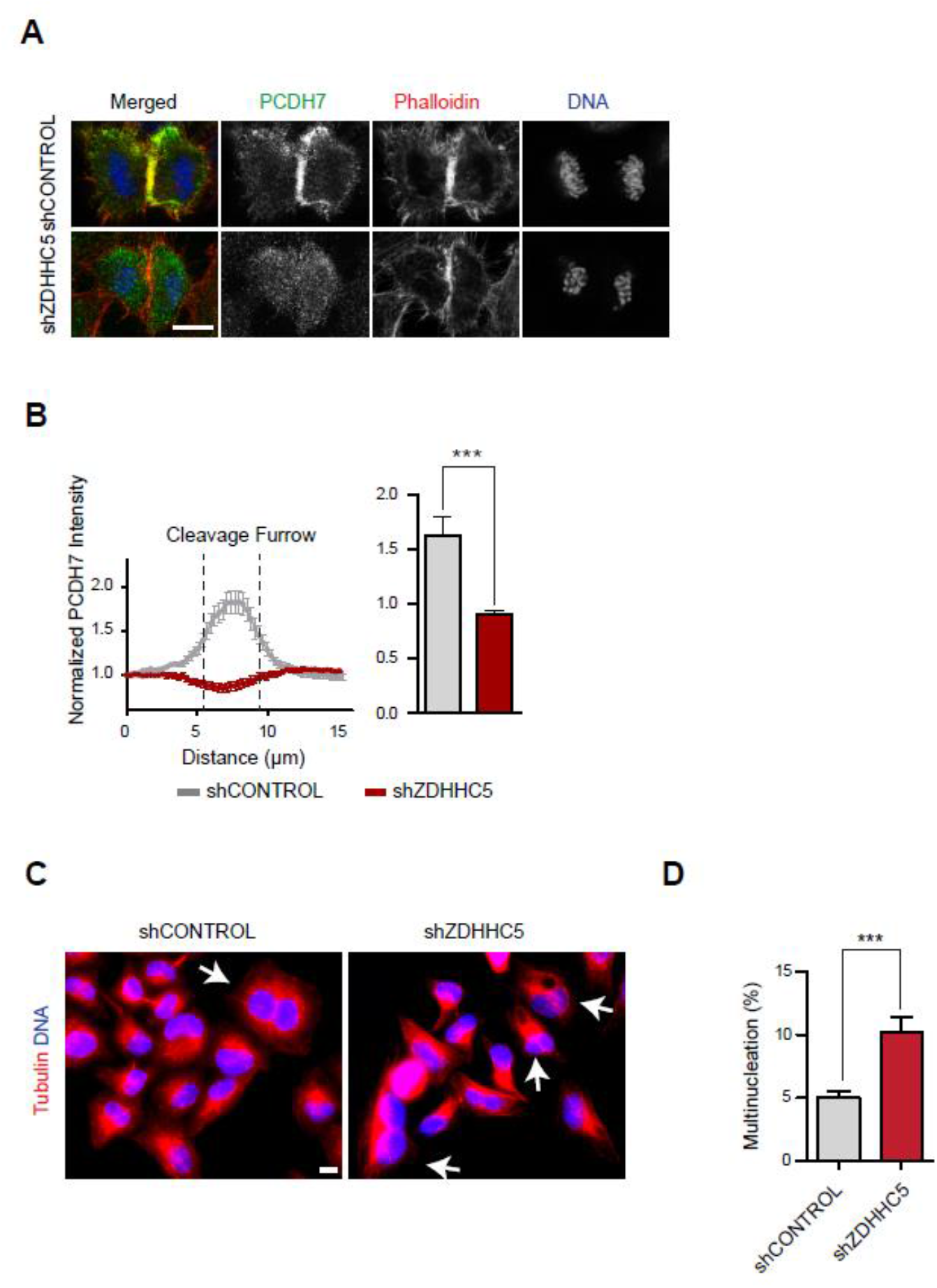
ZDHHC5 targets PCDH7 to the cleavage furrow and its expression contributes to cytokinesis. **A.** Representative maximum intensity projections of Z-stacks show PCDH7 (green, anti-GFP) localization at the cleavage furrow during cytokinesis in shCONTROL (top) and shZDHHC5 (bottom) treated PCDH7-GFP-BAC cells. Actin filaments (Phalloidin) are shown in red and DNA (DAPI) in blue. **B.** Intensity profiles of PCDH7 in the control (shCONTROL-gray) (n=35) and ZDHHC5 KD (shZDHHC5-red) (n=43) cells (left). The sum of PCDH7 intensities in between the dashed lines covering the +/- 15% distance around the middle zone, were quantified (right). Intensity profiles were obtained in ImageJ software for the indicated region of interest as previously described (Uretmen Kagiali et al., 2020). Statistics used *unpaired two-tailed t-test*, error bars: SD. **C.** Representative images of the multinucleated cells in control (shCONTROL) and ZSHHC5 knock-down (shZDHHC5) cells. Arrows indicate the multinucleated cells. **D.** Quantification of multinucleation percentages of control (shCONTROL) (n=1415, gray) and ZDHHC5 knockdown (shZDHHC5) (n=1276, red) cells. Statistics used *t test*, error bars:SEM. Scale bars: 10 µm. ***: p<0.001.

As ZDHHC5 localizes to the cleavage furrow together with PCDH7, next we examined whether ZDHHC5 depletion has any effect on cytokinesis. Intriguingly, we observed significantly increased multinucleation rate (8.5%) in shZDHHC5 cells in comparison to cells treated with control shRNA (4.6%) (**Figure 7C-D**). To draw a conclusion, ZDHHC5 localizes to the cleavage furrow, and targets PCDH7 to the cleavage furrow.

### Lack of PCDH7 at the cleavage furrow diminishes phospho Myosin Levels of Contractile Ring

To gain more insight to the cytokinesis defect of PCDH7 depleted cells, we monitored active RhoA and myosin levels at the cleavage furrow. Control and PCDH7 KO cells were transfected with a eGFP:RhoA Biosensor (Piekny and Glotzer, 2008) which binds only to active form of the RhoA thus, enables to track the RhoA activity. Live cell imaging was performed to compare the RhoA activity during cytokinesis in control (**Figure 8A**-top panel, **VideoS6**) and PCDH7 KO (**Figure 8A**-bottom panel, **VideoS7**) cells. Active RhoA levels at the cleavage furrow were significantly decreased in PCDH7 KO cells when compared with the control cells (**Figure 8B**, S5). Next, we examined active myosin levels in PCDH7 depleted cells by using anti-phospho-myosin II (S19) (Matsumura et al., 1998) (**Figure 8C**). The phosphorylated myosin is significantly enriched at the cleavage furrow in control cells (**Figure 8D**, gray). In PCDH7 KO cells, phospho-myosin intensity is significantly reduced at the cleavage furrow (**Figure 8D**, red). The expression of ectopic PCDH7 in PCDH7 KO cells significantly replenished the phospho-myosin levels (**Figure 8D**, blue).

**Figure 8.**
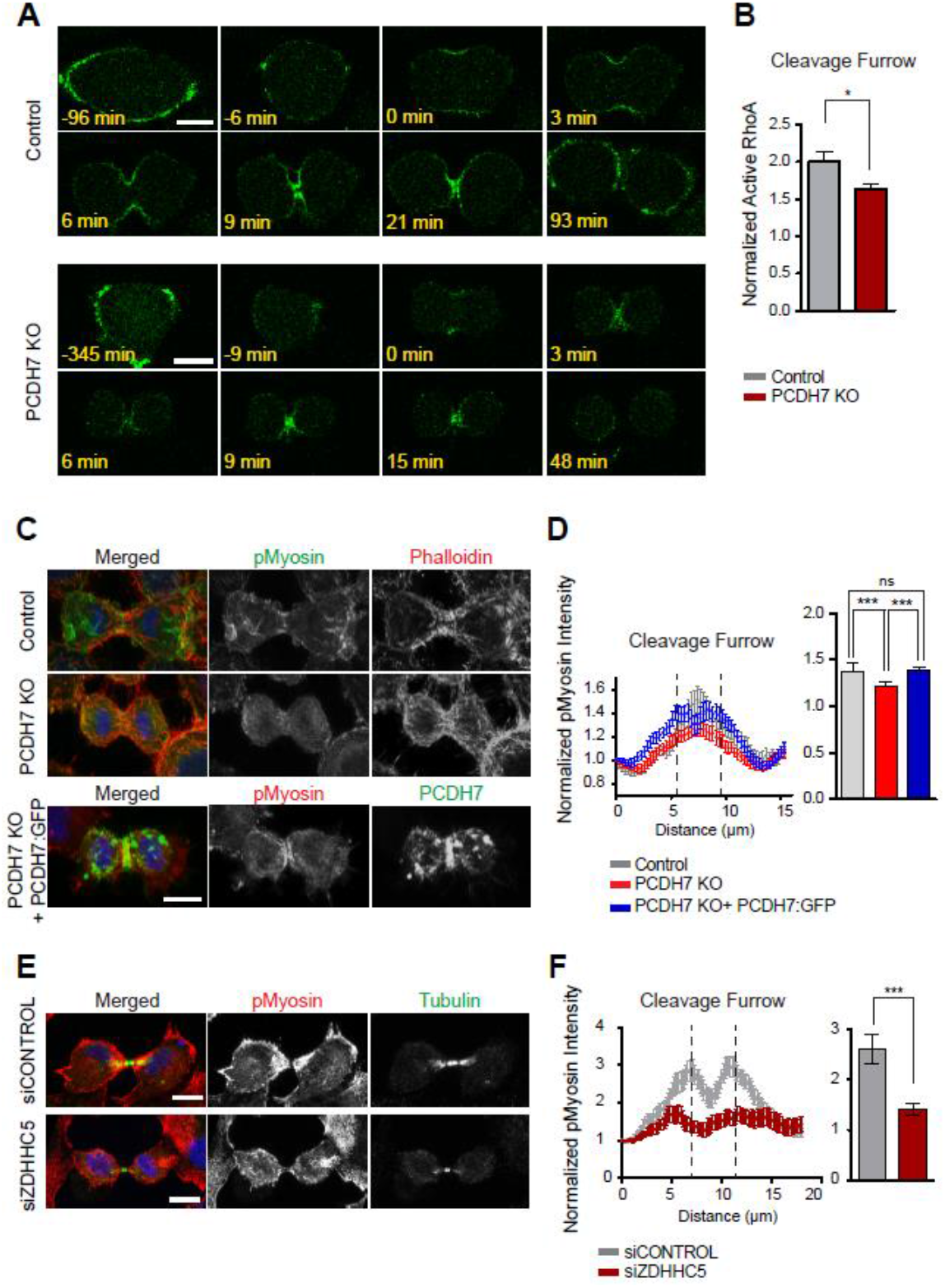
Active RhoA and Myosin phosphorylation levels are diminished at the cleavage furrow in PCDH7 depleted cells. **A.** Live imaging snapshots of control and PCDH7 knockout (PCDH7 KO) cells that are transfected with a pEGFP:RhoA-Biosensor vector to track active RhoA. Relative timing according to cleavage furrow ingression is shown in minutes. **B.** Quantification of the active RhoA levels at the cleavage furrow of control (n=44, gray) and PCDH7 KO (n=48, red) cells. The detail of the quantification method is illustrated in Figure S5. Statistics used *t test*, error bars:SEM. **C.** Representative fluorescence images displaying phospho-myosin II (S19) (green), actin filaments (red, Phalloidin), and DNA (blue, DAPI) localization in control (top) and PCDH7 KO (middle) cell. Rescue cells (PCDH7 KO+ PCDH7:GFP, bottom) display PCDH7:GFP (green), phospho-myosin II (S19) (red) and DNA (blue, DAPI). Maximum intensity projections of Z stacks are shown. **D.** Intensity profiles of pMyosin levels at the cleavage furrow in the control (gray) (n=24), PCDH7 KO (red) (n=24) and rescue (PCDH7 KO+ PCDH7:GFP) (blue) (n=41) cells (left). The sum of pMyosin intensities in between the dashed lines covering the +/- 15% distance around the middle zone, were quantified (right). Statistics used *one-way ANOVA with Benferonnis multiple comparison test*, error bars: SD **E.** Representative fluorescence images displaying phospho-myosin II (S19) (red), microtubules (green, anti-β-Tubulin) and DNA (blue, DAPI) localization in control (top panel) and ZDHHC5 knockdown (bottom panel) cells at cytokinesis stage. Maximum intensity projections of Z stacks are shown. **F.** Intensity profiles of pMyosin levels at the cleavage furrow in the control (gray) (n=28) and ZDHHC5 knock-down (red) (n=23) cells (left). The sum of pMyosin intensities in between the dashed lines covering the +/- 15% distance around the middle zone, were quantified (right). Statistics used *unpaired two-tailed t-test*, error bars: SD. Scale bars: 10 µm, *: p<0.05; **: p<0.01; ***: p<0.001.

In agreement with ZDHHC5 dependency of PCDH7, similar to PCDH7, depletion of ZDHHC5 also caused a significant decrease at the phospho-myosin (S19) levels at the cleavage furrow (**Figure 8 E, F**). Taken together, our data suggest that ZDHHC5-PCDH7 axis positively regulates RhoA and myosin activity at the cleavage furrow, thus contributes to the fidelity of cytokinesis.

## DISCUSSION

This study aimed to understand the function and the underlying mechanism of cell cycle dependent localization of PCDH7. Our analysis revealed an unprecedented role of palmitoylation in the translocation of PCDH7 to the mitotic cell surface and cleavage furrow which promotes myosin phosphorylation at the cleavage furrow. Palmitoylation is a reversible posttranslational modification that plays a key role in controlling protein targeting by increasing the hydrophobicity of a protein (Linder and Deschenes, 2007). Our data support a mechanism by which palmitoylation of PCDH7 stabilizes its cell surface and cleavage furrow localization. Indeed, the cell surface and cleavage furrow localization during mitosis and cytokinesis respectively, was dependent on the palmitoylation, whereas the localization to the cell-to-cell contacts during interphase was not affected by the palmitoylation inhibition. Our FRAP analysis revealed that the turnover rate and dynamics of molecules at the cell-cell contact region and cell surface are different, the latter is faster. It is possible that trans interactions are more stable and palmitoylation is dispensable or less critical for trans clustered molecules at the cell-cell contact regions. Our analysis revealed that PCDH7 gets palmitoylated however our attempts of finding palmitoylation sites by mutating some potential cysteine residues that affect protein localization failed. It is rather challenging to determine the palmitoylation sites because of the lack of a consensus motif. Besides cysteine, palmitoylation can also occur on serine and lysine residues. In addition, a cumulative effect of palmitoylation on multiple cysteines, and even on serine or lysine (Brownlee and Heald, 2019) residues might be involved, thus mutating individual residues might not be enough to mimic the unpalmitoylated form. Similarly, PCDH1, which shares 46% homology with PCDH7, was also stated to be palmitoylated and localize to the membrane in a palmitoylation-dependent manner. However, mutating one palmitoylated cysteine residue was not enough to mimic the palmitoylation inhibitor and exhibit the membrane localization phenotype (Kahr et al., 2013). More work is required to understand the molecular details in palmitoylation of non-clustered protocadherins.

Our BioID based proximity interactome identified a palmitoyl transferase ZDHHC5, as a significant interactor of PCDH7. In contrast to the majority of PAT members that localize to Golgi or endoplasmic reticulum, ZDHHC5 localizes to the plasma membrane (Ohno et al., 2006). ZDHHC5 has been implicated in multiple cellular processes such as endocytosis, cell adhesion, Na-pump activity, and pathogen-host interaction partly by regulating the localization of related proteins (Plain et al., 2020; Pradhan et al., 2021; Woodley and Collins, 2019, 2021). In this study for the first time, we analyzed the function of ZDHHC5 in the context of cell division. At metaphase, ZDHHC5 localizes to the mitotic cell surface and retraction fibers, and it concentrates at the cleavage furrow during cytokinesis. The translocation of PCDH7 to the mitotic cortex and cleavage furrow occurs in ZDHHC5 palmitoylation activity dependent manner.

Loss of ZDHHC5 caused cytokinesis defects and statistically significant increase in the rate of multinucleated cells. Imaging of protein fatty acylation in cells undergoing cell division revealed that at metaphase S-palmitoylation enriches at the cell surface and around the spindle, as cells progress into cytokinesis, palmitoylated proteins are concentrated at the cleavage furrow (Hannoush and Arenas-Ramirez, 2009). Based on those findings, it is entirely possible that PATs and partly ZDHHC5 may have a wider role in organizing the mitotic cell cortex and targeting cell division-related proteins to the plasma membrane and cleavage furrow. PCDH7 might be the very first example of many of cell cycle dependent palmitoylated proteins. Although many studies have indicated the role of protein palmitoylation in protein trafficking (Linder and Deschenes, 2007) and different intracellular signaling pathways (Resh, 2006), the importance of palmitoylation in the context of cell division has been emerging only recently. Depalmitoylation activity has been shown to be required for the unequal partitioning of the Notch and Wnt signaling during asymmetric cell division (Stypulkowski et al., 2018). Palmitoylation-dependent membrane association of importin α has been shown to affect the mitotic spindle and nuclear scaling during Xenopus embryogenesis (Brownlee and Heald, 2019). Our findings attribute new functional aspects to the role of palmitoylation in cell division. Future studies will expand the palmitoylation dependent regulatory mechanisms during cell division by investigating palmitoyltransferases and their target molecules that function in mitosis and cytokinesis.

What is the function of PCDH7 during cell division? Our previous study showed that PCDH7 is required for the development of full mitotic rounding pressure (Ozlu et al., 2015). Rounding pressure is dependent on myosin II activity (Stewart et al., 2011a; Stewart et al., 2011b). In the present study, we have shown that the knockout of PCDH7 caused a statistically significant increase in cytokinesis failure in HeLa cells. The multinucleation phenotype was moderate but significant. In our previous study, although we observed a reduced rounding pressure in PCDH7 siRNA cells, the cells were still able to round up. The weakness of the observed phenotypes may be due to a functional redundancy between PCDH1 and PCDH7 and the fact that multiple parallel pathways act together for mitotic rounding and cytokinesis (Cramer et al., 1994; Ramkumar and Baum, 2016).

Actomyosin network is important for mitotic rounding and cleavage furrow formation and both processes seem to be affected in the absence of PCDH7. A previous genome-wide RNAi screen also reported binucleation and cell migration defects in PCDH7 depleted cells (Neumann et al., 2010). Indeed, we observed that active Rho A and myosin levels at the cleavage furrow were significantly reduced in the absence of PCDH7 when compared to the control cells. Previously it was reported that PCDH7 increases phospho-myosin light chain levels to enhance anchorage-independent cell growth (Wang et al., 2020). Similarly, in a parallel study, we observed that PCDH7 overexpression enhances phospho-myosin levels during cell migration in Retinal Pigment Epithelial (RPE) cells and PCDH7 interacts with PP1cβ, the catalytic subunit, and MYPT1, the myosin targeting subunit of myosin phosphatase holoenzyme (Qureshi et al., 2021). Previous studies also reported that PCDH7 behaves as a signaling molecule more than an adhesion molecule and inhibits phosphatase activities of PP1a and PP2A (Wang et al., 2020; Zhou et al., 2017). It is possible that PCDH7 regulates phosphorylation level of myosin through its interaction with phosphatases or Rho-ROCK modulators. Recently, it has been reported that PCDH7 functions in osteoclast differentiation by regulating RhoA and Rac1. Loss of PCDH7 expression impairs RhoA activation in pre-osteoclasts (Kim et al., 2021). In agreement with those studies, we also observed that depletion of PCDH7 impairs active RhoA and myosin levels at the cleavage furrow. Further study is necessary to elucidate the molecular details of PCDH7 dependent regulation of RhoA and myosin activity during cytokinesis.

In summary, our study suggests a new pathway for the cell cycle dependent protein localization during cell division. We suggest that the ZDHHC5 dependent targeting of PCDH7 to the cell cortex and cleavage furrow contributes to cortical remodeling. The molecular details of palmitoylation dependent cortical regulation during cell division will unravel the fundamental redundant pathways existing in human cells.

## MATERIALS AND METHODS

### Cell Lines and Culture

HeLa S3 cells (ATCC CCL-2.2, female) and HEK293T cells (a kind gift from Dr. Tamer Önder) were grown in Dulbecco’s modified Eagle’s medium (DMEM) (Sigma-Aldrich, D6429) supplemented with 1% penicillin-streptomycin (P/S) (Capricorn, PS-B) and 10% fetal bovine serum (FBS) (Gibco, 10270106). PCDH7-GFP-BAC (MCP Ky 5914 T457) (Poser et al., 2008) transgenic cell line was a kind gift from Dr. Ina Poser and was grown in DMEM supplemented with 1% P/S, 10% FBS, and 400 μg/ml G418 (Santa Cruz, sc-29065A).

To inhibit the palmitoylation, cells were incubated with 100 µM 2-Bromopalmitate (2BP) (Sigma, 21604) overnight (Webb et al., 2000).

### Cell Synchronization

Cells synchronization to interphase, mitosis (Ozlu et al., 2010) and cytokinesis (Hu et al., 2008; Karayel et al., 2018) was performed as previously described. Cells were synchronized to interphase by double thymidine block using 2 mM thymidine (Santa Cruz, sc-296542). Subsequently, for monopolar mitosis, cells were treated with 10 µM S-trityl-L-cysteine (STC) (Sigma-Aldrich, 164739). For the monopolar cytokinesis, cells were treated with 10 µM STC and then 100 µM purvalanol A (Tocris Bioscience, 1580). For bipolar synchronization, cells were incubated with 10 ng/ml of nocodazole (Calbiochem, 487928) for 5 hours for mitosis and released from nocodazole for 1 hour for cytokinesis.

### Immunostaining and Microscopy

For immunostaining, cells were plated on coverslips (12 mm), fixed with 3% paraformaldehyde (PFA), blocked, and incubated with primary and secondary antibodies in 2% BSA in PBS containing 0.1% Triton-X. The following antibodies and reagents were used: Beta-Tubulin (Cell Signaling, CS2128S), Alpha Tubulin (Cell Signaling, 3873S), Tubulin (Abcam, ab6160), GFP (non-commercial/Invitrogen, A11120), HA (Abcam, ab16918), pMyosin (Cell Signaling, CS3675), ZDHHC5 (Atlas Antibodies, HPA014670). Alexa Fluor®-488 and Alexa Fluor®-555 (Cell Signaling), Streptavidin-Alexa Fluor 488 (Invitrogen, S32354), Phalloidin iFlour (Abcam, ab176756), DAPI (Sigma Aldrich, D8417).

For live imaging, ibiTreat, ibidi μ-Slide 8 Well (ibidi, 80826) or μ-Dish 35 mm (ibidi, 81156) plates were used.

Confocal microscopy was performed in Leica DMi8/SP8 TCS-DLS (LAS X Software) laser scanning confocal microscope using 40× Plan Apo 1.3 NA and 63x Plan Apo 1.4 NA oil-immersion objectives.

Live imaging experiments were performed using Leica DMi8 widefield fluorescence microscope (LAS X Software) or Leica DMi8/SP8 TCS-DLS confocal microscope equipped with 37°C and 5% CO_2_ chambers. 63× Plan Apo 1.4 NA oil immersion objective or 20× PL FLUOTAR L 0.40 NA objectives were used. Single images or Z-stacks were acquired every 3 minutes and a single focal plane was used in the figures unless specified in the figure legends.

Apart from FRAP data, all images were analyzed in Fiji. Graphs and statistical data were generated in GraphPad Prism. Statistical details of each experiment including the statistical test used, the exact value of n, and definition error bars can be found in the figure legend for each figure.

### Fluorescence Recovery After Photobleaching (FRAP) Analysis

#### Microscopy setup

The microscopy set-up included a frequency-doubled femtosecond-pulsed Ti:Sa solid-state tunable laser source (Chameleon Ultra II, Coherent) equipped with a second harmonic generator. Laser output was tuned to 488 nm and the beam was directed through mirrors and a Keplerian telescope to the inverted microscope (Eclipse TE2000-U; Nikon) equipped with a dichroic mirror (Chroma, Q495LP) and 60X oil-immersion objective (Nikon Apo TIRF, NA=1.49). A 300 mm focal length lens was placed right before the microscope to focus the laser at the back focal plane of the microscope objective to obtain wide-field illumination. The microscope was equipped with two different cameras for brightfield and fluorescence image acquisitions. Brightfield images were captured by a CCD camera (Thorlabs, DCU223M). Fluorescence images were captured by an EMCCD camera (Hamamatsu ImagEM C9100-13) placed after an emission filter with a pass band 530 ± 30 nm. Photobleaching was performed by removing the neutral density optical filter and the lens right before the microscope, and simultaneously focusing the laser light at the desired area for 1 second. FRAP image acquisition was done every five seconds using an automated shutter and a minimum of 130 frames were captured following photobleaching of each sample.

#### Data Processing

Image analysis was done by using a MATLAB code based on the double normalization algorithm (Phair et al., 2004). In this method, the overall decrease in the fluorescence intensity of the samples is also considered and the normalized intensity of the region of interest (ROI). *I*_*normalized*_ is given by:

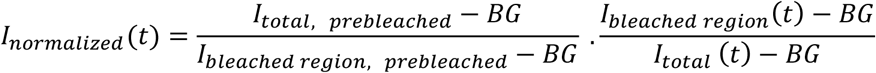

In which, *I*_*total, prebleached*_ and *I*_*bleached region, prebleached*_ are prebleached intensities of the whole cell and the ROI, respectively. *I*_*bleached region*_*(t)* and *I*_*total*_ *(t)* are corresponding intensities at time *t. BG* is the intensity of the background region. The total number of frames and the first post bleached frame was defined for each sample. The size of the selected background and the ROI areas in each sample was fixed to 14×14 pixels (Kappel, 2004). Unhealthy cells and the motile cells that obstruct the tracking of the ROI throughout the frames are discharged from the data. Obtained *I*_*normalized*_*(t)* data for each cell then is fitted to 1 − *Ae*^−*at*^ − *Be*^−*bt*^ (Phair et al., 2004) to get half time values *t*_1/2_. The recovery percentage was also calculated for each cell. The *Mann–Whitney U test* was applied to reveal statistical differences in the recovery and the half-time between samples. Average recovery curves for each group of cells (cell-cell contacts in interphase and mitotic plasma membrane) were obtained by normalizing all individual recovery graphs recorded within a group to 1 and calculating the average points as well as standard deviations of the recovery data at specified frames (time instances) after photobleaching.

### Transfection and Viral Transduction

Cells were transfected using polyethylenimine (PEI) or Lipofectamine 2000 protocols (Invitrogen, 11668019). For PEI protocol, transfection mixtures were prepared in Opti-MEM reduced serum medium (Invitrogen,31985047) using PEI with 60 μg DNA/150mm dishes in the 3:1 ratio. After 30 minutes of incubation at room temperature, the transfection mixture was added onto cells. For Lipofectamine 2000 protocol, the manufacturer’s instructions were followed.

Murine ZDHHC5 and its catalytically inactive mutant ZDHHC5 (C134S) plasmids were kindly provided by Prof. William Fuller, Institute of Cardiovascular and Medical Sciences, College of Medical, Veterinary and Life Sciences, University of Glasgow, Glasgow, UK. pEGFP-RhoA Biosensor plasmid was kindly provided by Micheal Glotzer (Addgene plasmid #68026).

For viral transduction, lentiviruses were packaged in HEK293T cells using PEI transfection of target sequence containing vector (pLenti), packaging vector (psPAX2, Addgene 12260), and envelope vector (pVSVg, Addgene 8454). Viral particles were collected after 48 and 72 hours of transfection and used to infect HeLa S3 in the presence of 2 μg/ml protamine sulfate (Sigma-Aldrich, P4505) as coadjutant. Transduced cells were then selected with the appropriate selective antibiotic.

### Identification of Proximity Dependent Interactions (BioID)

PCDH7 sequence was amplified from PCDH7:eGFPN1 using specific primers: 5’-GTC AGC TAG CAC CAT GCT GAG GAT GCG GACC-3’, 5’-GCT AGA ATT CGC CCT CCC TGG GAT ATT TAA ATA TAT TTG-3’ and cloned into BioID vector (pcDNA3.1 MCS-BirA* (R118G) Addgene; 36047).

Proximity-dependent biotinylation was performed as previously described (Roux et al., 2012). Briefly, cells were transfected with BioID vector and incubated with 50 μM Biotin (Invitrogen, B20656) during interphase and mitosis synchronization.

Cell pellets were lysed in lysis buffer (50 mM Tris, pH 7.4; 500 mM NaCl; 0.4% SDS; 5 mM EDTA; 2% TritonX; 1 mM DTT; Protease Inhibitor) and incubated with Streptavidin beads (Pierce,53117) overnight at 4°C in a tube rotator. Fractions of the whole-cell lysate (WCL) and Unbound (Ub) were kept at -20°C for further analysis. Beads were washed with Wash Buffer 1 (2% SDS in dH_2_O), Wash Buffer 2 (2% deoxycholate; 1% TritonX; 50 mM NaCl; 50 mM Hepes, pH 7.5; 1 mM EDTA), Wash Buffer 3 (0.5% NP-40; 0.5% deoxycholate; 1% TritonX; 500 mM NaCl; 1 mM EDTA; 10 mM Tris, pH 8.1) and Wash Buffer 4 (50 mM Tris, pH 7.4; 50 mM NaCl). For Western blot analysis, bound proteins were eluted from the streptavidin beads with 50 µl of Laemmli-DTT sample buffer containing 500 nM D-Biotin at 98°C by shaking at 1,000 rpm in a ThermoMixer (Eppendorf™). For mass spectrometry analysis, on-bead tryptic digestion was performed. Briefly, beads were washed with urea buffer (8 M urea (Sigma-Aldrich, A2383) 0.1 M Tris/HCl pH 8.5). Then, bead-bounded proteins were reduced with 100 mM Dithiothreitol (DTT) (Sigma-Aldrich, 43815) and alkylated using 100 mM iodoacetamide (Applichem, A1666). After alkylation, beads were washed with 50 mM ammonium bicarbonate (Applichem, A3583) and incubated with Trypsin (Thermo Scientific,25247) at 37°C overnight (14-16 hours) in ThermoMixer (Eppendorf™), the resulting digested peptides were collected and desalted using C18 STAGE Tips. The experiment was performed in 4 biological replicates with a minimum of 2 technical replicates for each.

### Mass Spectrometry and Data Analysis

Peptides were analyzed by online C18 nanoflow reversed-phase nLC (NanoLC-II, Thermo Scientific) or C18 nanoflow reversed-phase HPLC (Dionex Ultimate 3000, 3500 RSLC nano, Thermo Scientific) connected with an orbitrap mass spectrometer (Q Exactive Orbitrap, Thermo Scientific). Samples were separated in an in-house packed 100 μm i.d. × 23 cm C18 column (Reprosil-Gold C18, 5 μm, 200 Å, Dr. Maisch) using 80-minutes linear gradients from 5-25%, 25-40%, 40-95% acetonitrile in 0.1% formic acid with 300 nL/min flow in 100 minutes total run time. The scan sequence began with an MS1 spectrum (Orbitrap analysis; resolution 70,000; mass range 400–1,500 *m*/*z*; automatic gain control (AGC) target 1e6; maximum injection time 32 ms). Up to 15 of the most intense ions per cycle were fragmented and analyzed in the orbitrap with Data Dependent Acquisition (DDA). MS2 analysis consisted of collision-induced dissociation (higher-energy collisional dissociation (HCD)) (resolution 17,500; AGC 1e6; normalized collision energy (NCE) 26; maximum injection time 85 ms). The isolation window for MS/MS was 2.0 *m*/*z*.

Raw files were processed with Proteome Discoverer 2.3 (Thermo Scientific) software. Carbamidomethylation of cysteine residues was used as fixed modification, and acetylation (protein N-termini) and oxidation of methionine residues were used as variable modifications. Maximal two missed cleavages were allowed for the tryptic peptides. The precursor mass tolerance was set to 10 ppm and fragment mass tolerance was set to 0.02 Da. Both peptide and protein false discovery rates (FDRs) were set to 0.01. The other parameters were used with default settings. The database search was performed against the human Uniprot database (release 2015) containing 21,039 entries using the SEQUEST HT search engine integrated into the Proteome Discoverer environment.

### Network Analysis

The spectral counts of proteins were used to calculate fold change ratios and FDR values for identified proteins using the qspec-param program of qprot_v1.3.5 (Choi et al., 2015). Proteins are filtered with a 0.05 cut-off for FDR values. Significant protein hits were loaded into the String database v11.0 (Szklarczyk et al., 2019) by Cytoscape StringApp (Doncheva et al., 2019) with 0.7 confidence. MCODE clustering of the network was performed by Cytoscape (version 3.7.2) and its plugin clustermaker (Cline et al., 2007). GO and KEGG enrichment analysis of the network was performed via g:Profiler (Raudvere et al., 2019).

### Proximity Ligation Assay (PLA)

Duolink PLA Kit (Sigma-Alrich, 92101) was used based on the manufacturer’s procedure. Briefly, HeLa S3 PCDH7-GFP-BAC cells were seeded onto glass coverslips and synchronized to cytokinesis with bipolar synchronization. The cells were fixed by using 3.2% PFA and permeabilized by 0.1 % TritonX-TBS. After blocking with Duolink blocking solution for 30 minutes at 37ºC, cells were incubated overnight at 4ºC with the corresponding pair of primary antibodies: anti-GFP (Invitrogen, A11120) for PCDH7 and anti-ZDHHC5 (Sigma-Alrich, HPA014670). “Only one primary antibody-treated” samples for each antibody were used as controls. The cells were washed and incubated with PLA probes for 1 hour at 37ºC. After another wash, cells were treated with the ligase for 30 minutes at 37ºC. The washing step is repeated, then the cells were incubated with the polymerase for 100 minutes at 37ºC. After final washes, the slides were mounted with a coverslip by using Duolink Mounting Medium with DAPI and incubated for 15 minutes before sealing.

### GFP-trap Pull-Down Assay for the Protein Interactions

HeLa S3 cells expressing GFP (emptyGFP) or PCDH7:GFP were arrested in mitosis and used for pulldown assay. The pellets were dissolved in PBS with 1% Triton X-100, EDTA-free protease inhibitor (Thermo Pierce, 88666), and Phostop (Roche, 4906845001) and then homogenized and centrifuged at 14,000 rpm for 10 minutes at 4ºC. Protein concentrations were determined by BCA protein assay (Pierce, 23225) and equal protein from each sample was loaded into pre-conditioned GFP-Trap A beads (gta-20; ChromoTek). Aliquots of the input sample(I) were saved for further analysis. After incubation for 3 hours at 4ºC, the unbound samples (Ub) were collected, GFP-Trap A beads were washed and proteins were eluted in 2x Laemmli with 100mM DTT by boiling for 10 minutes at 95ºC

### Western Blotting Analysis

Samples were separated by molecular weight using 10% SDS-PAGE gels and transferred to a nitrocellulose membrane. The membrane was blocked with 4% w/v nonfat dry milk in TBS-0.1% Tween-20 and probed with 1 μg/ml of the described primary antibody diluted in 2% BSA TBS-0.1% Tween-20. The signal was visualized using ECL (Pierce, 32106) detection of the HRP-conjugated secondary antibodies (Cell Signaling, 7074S, 70765). The following primary antibodies were used in the Western blot analysis: PCDH7 (Abcam, ab139274), EGFR (Santa Cruz, SC-03), Tubulin (Cell Signaling, 3873), Actin (Abcam, ab6276), Phospho-Histone H3 (Upstate; 06-570) and Biotin (non-commercial), GFP (non-commercial), ZDHHC5 (Atlas Antibodies, HPA014670), Calnexin (Abcam, ab22595).

### Detection of Palmitoylation by Acyl-Biotin Exchange Assay

For detection of the protein palmitoylation, the Acyl-Biotin Exchange (ABE) procedure was performed as described (Wan et al., 2007). Briefly, cell pellets were lysed in ice-cold Lysis buffer (LB; 150 mM NaCl, 50 mM Tris, 5 mM EDTA, pH 7.4 with 10 mM NEM, 2x PI, and 2x PMSF). After homogenization, membrane proteins were enriched by using high-speed centrifugation (Optima MAX-XP Ultracentrifuge, TLA-120.2 rotor) at 200,000g for 30 min at 4ºC. The membrane enriched pellet was dissolved in the LB with 10 mM NEM, 1x PI and 1x PMSF, and 1.7% Triton X-100 and incubated at 4ºC for 2 hours. To remove particulates, the sample was centrifuged at 250g for 5 minutes at 4ºC. Chloroform-methanol (CM) precipitation was applied to precipitate the proteins. The pellet was air-dried for 2-3 minutes and 4% SDS buffer (4SB; 4% SDS, 50 mM Tris, 5 mM EDTA, pH 7.4) with 10 mM NEM was added into the sample and incubated for 20 minutes at 37ºC to dissolve the protein pellet completely. After NEM incubation overnight at 4ºC (1 mM NEM, 1x PI, 1 mM PMSF, and 0.2% Triton X-100), three sequential CM precipitations were applied to remove NEM from the sample. After the final CM precipitation, the pellet was dissolved in 4SB and sample was divided into two equally as HA- and HA+ sample and incubated for 1 hour at room temperature with HA- (50 mM Tris, 1 mM HPDP–biotin, 0.2% Triton X-100, 1 mM PMSF, 13 PI, pH7.4) and HA+ (0.7 M hydroxylamine, 1 mM HPDP–biotin, 0.2% Triton X-100, 1 mM PMSF, 1xPI pH 7.4) buffers respectively. Then, proteins were precipitated by CM precipitation and dissolved pellets were incubated in low HPDP–biotin buffer (150 mM NaCl, 50 mM Tris, 5 mM EDTA, 0.2 mM HPDP–biotin, 0.2% Triton X-100, 1 mM PMSF, 1x PI, pH 7.4) for 1 hour at room temperature. Three sequential CM precipitation was performed to remove unreacted biotin. The protein pellets were dissolved in 4SB and then SDS was diluted to 0.1 % by addition of 0.2% Triton X-100, 1x PI, 1mM PMSF, and samples were incubated at room temperature for 30 minutes. Then, samples were loaded on pre-conditioned Streptavidin Plus UltraLink Resin (Pierce, 53117) and incubated for 90 minutes at room temperature. Unbound fractions (Ub) from both samples were saved, beads were washed with LB containing 0.1%SDS and 0.2% Triton X-100 three times, and bound proteins were eluted in 2x Laemmli with 1% β-mercaptoethanol by boiling for 10 minutes at 95ºC.

### Triton X-114 Extraction

Hydrophobic proteins were extracted from the hydrophilic ones using the Triton X-114 (TX-114) extraction protocol as described (Bordier, 1981; Taguchi et al., 2013). Briefly, precondensation of the TX-114 was performed by repeated cycles of clarifying at 4°C and incubation at 37°C to separate the detergent phase. Cells were lysed using 2% TX-114 lysis buffer in PBS and lysate was cleared by centrifugation at 16,100 g for 3 minutes at 4°C. The lysate was then centrifuged at 22,000 g for 10 minutes at room temperature for phase separation. The aqueous phase was removed, the detergent phase was washed with 0.1% TX-114 wash buffer and clarified on ice and incubated at 37°C for phase separation. Centrifugation and washing steps were repeated 2 more times and detergent and aqueous phases were collected for further analysis.

### Surface Labelling and Pulldown of Cell Surface Proteins

Plasma membrane proteins were labeled with sulfo-NHS-SS-biotin for membrane enrichment as previously described (Özkan Küçük et al., 2018). Briefly, cells were incubated with 5 mM S-NHS-SS-biotin (Pierce, 21331) for 30 minutes at 4°C with gentle shaking, the reaction was quenched with glycine, and cells were snap-frozen. Cells were lysed in a buffer (10 mM TrisCl pH 7.6, 0.5% SDS, 2% NP40, 150 mM NaCl, 1 mM EDTA, 10 mM Iodoacetamide) supplemented with protease inhibitors (Pierce, 88666) and lysates were incubated with pre-conditioned Streptavidin Plus UltraLink Resin (Pierce, 53117) overnight at 4°C. Unbound samples were collected and beads were washed with lysis buffer three times. Biotinylated surface proteins were eluted by boiling at 70°C for 20 minutes in SDS sample buffer including 100 mM DTT with agitation. All fractions including whole cell lysate (Input-I), unbound (U), and plasma membrane enriched (Elute-E) were analyzed by Western blot.

### RNAi Mediated Gene Silencing

ZDHHC5 expression was knocked down using shRNA and siRNA approaches. shZDHHC5 was gifted from Dr. G. Ekin Atilla-Gökcümen (Pradhan et al., 2021) and pLKO.1 was gifted from Dr. Elif Nur Fırat Karalar. The lentiviral particles of shZDHHC5 (target sequence 5’CCCAGTTACTAACTACGGAAA3’ in pLKO.1 vector) and empty pLKO.1 (Addgene, 8453) were packaged in HEK293T cells and used to transduce PCDH7-GFP-BAC cells. Virus incorporated stable cells that stably express shRNA’s were obtained after puromycin selection.

siGENOME siRNA pools that target ZDHHC5 (siZDHHC5, Dharmacon, D-026577-01-0020, target sequence: GGACUAAGCCUGUAUGUGU) and Non-targeting siRNA (Dharmacon, D-001210-01-05) were transfected with Lipofectamine RNAiMAX Transfection Reagent (Thermo Scientific) according to manufacturer’s instructions. Briefly, cells were seeded and transfected with 15pmol of siRNA two times, at 24 hours and 48 hours. After 72 hours after seeding, cells were either pelleted for Western blot analysis or fixed for immunofluorescence analysis.

### CRISPR Based Knock Out

Single guide RNAs (sgRNAs) that target PCDH7 were designed using online “CRISPR design toll” (http://crispr.mit.edu). The following oligonucleotide sequences were used as top/bottom pairs: sg1 5’-CACCGCGACGTCCGCATCGGCAACG-3’/5’-AAACCGTTGCCGATGCGGACGTCG-3’, sg5 5’-CACCGCATCGTGACCGGATCGGGTG-3’/5’-AAACCACCCGATCCGGTCACGATG-3’, sg6 5’-CACCGCGGGCTTCTCTTTGGCGCGC-3’/5’-AAACGCGCGCCAAAGAGAAGCCCGC-3’. All sgRNAs were cloned into lentiCRISPR plasmid (Shalem et al., 2014) (pXPR_001, 49535, Addgene) as described in (Ran et al., 2013).

PCDH7 knockout cell lines were generated by lipofectamine transfection of CRISPR plasmid to HeLa S3 cells followed by antibiotic selection. Single colonies were isolated with serial dilution of the pool population and PCDH7 knockout clones were selected after verifying the absence of PCDH7 protein expression with Western blot. CRISPR rescue cell lines were generated by viral transduction of pLenti (Campeau et al., 2009) PCDH7:eGFP plasmid to PCDH7 knockout cells.

## AUTHOR CONTRIBUTIONS

N.E.Ö. designed and performed the experiments, analyzed the data, and wrote the manuscript. B.N.Y. and B.S.D performed experiments together with N.E.Ö. M.H.Q. designed and created CRISPR knockout cell lines and PCDH7:GFP constructs. A.Ka. analyzed the BioID data. N.B. performed FRAP experiments together with N.E.Ö and analyzed the FRAP data. A.Ki. designed and analyzed FRAP experiments. N.Ö. designed the research, provided funding and wrote the manuscript.

## ACKNOWLEDGMENTS

This study is funded by TUBITAK 1001 (116Z305) to N.Ö. We gratefully acknowledge Büşra Aytül Akarlar and the Proteomics Facility of Koç University (KUPAM) for the technical assistance in mass spectrometry analyses. We thank Dr. Bilal Ersan Kerman for the critical discussions throughout the project. We also thank Dr. Alexandr Jonas for the critical discussions and interpretations of FRAP data. We thank Dr. Timothy Mitchison for the anti-biotin antibody and the critical reading of the manuscript. We thank Dr. G. Ekin Atilla-Gökcümen for sharing ZDHHC5 shRNAs, Dr. Elif Nur Fırat Karalar for sharing pLKO.1 plasmid and anti-GFP antibody, and Dr. Tamer Önder for sharing HEK293T cells and Dr. William Fuller for sharing murine ZDHHC5 plasmids. We also thank Dr. Nazan Saner and Dr.Aydanur Şentürk for the critical reading of the manuscript. We gratefully acknowledge the permission to use the facilities of the Cellular and Molecular Imaging Core of Koç University Research Center for Translational Medicine funded by the Republic of Turkey Ministry of Development.

## CONTACT FOR REAGENT AND RESOURCE SHARING

Further information and requests for resources and reagents should be directed to and will be fulfilled by the Lead Contact, Nurhan Özlü.

All unique/stable reagents and codes generated in this study are available from the Lead Contact without restriction.

## LIST OF SUPPLEMENTAL ITEMS

**Figure S1**. Related to Figure 1 & Figure 2.

**Figure S2**. Related to Figure 3.

**Figure S3**. Related to Figure 4, Figure 5 & Figure 6, Figure S4.

**Figure S4**. Related to Figure 6.

**Figure S5**. Related to Figure 8.

**Video S1**. Related to Figure 1. Live imaging video of HeLa S3-PCDH7:GFP cells during cell division.

**Video S2**. Related to Figure 2. Live imaging of HeLa sgNT LifeAct RFP control cells during cell division.

**Video S3**. Related to Figure 2. Live imaging of HeLa PCDH7 KO LifeAct RFP cells during cell division.

**Video S4**. Related to Figure S3. Live imaging video of PCDH7-GFP-BAC expressing control cells (DMSO treated).

**Video S5**. Related to Figure S3. Live imaging video of PCDH7-GFP-BAC expressing cells treated with palmitoylation inhibitor 2BP.

**Video S6**. Related to Figure 8. Live imaging video of HeLa sgNT control cells expressing eGFP:RhoA-Biosensor during cell division.

**Video S7**. Related to Figure 8. Live imaging video of HeLa PCDH7 knockout cells expressing eGFP:RhoA-Biosensor during cell division.

**Table S1**. Related to Figure 3A. List of PCDH7 proximal interactome.

## DATA AVAILABILITY

“The mass spectrometry proteomics data have been deposited to the ProteomeXchange Consortium via the PRIDE (Perez-Riverol et al., 2019) partner repository with the dataset identifier PXD029246”.

## Supplementary Figures

**Figure S1.**
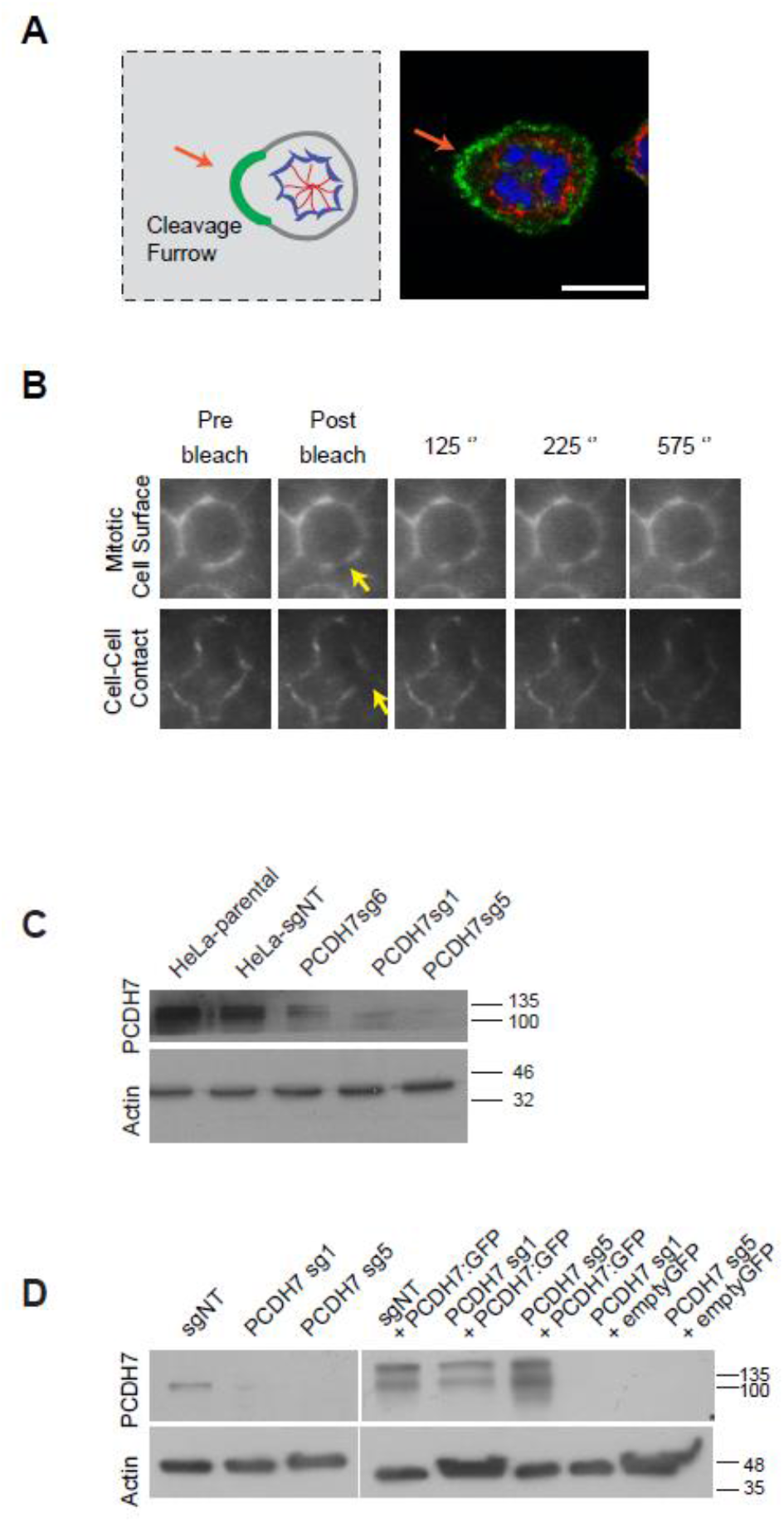
Characterization of cellular localization of PCDH7 and knockout cells. **A.** Illustration (left) and a representative fluorescence image (right) of monopolar cytokinesis cells. Green line and orange arrow marks bud-like structure that corresponds to the monopolar cleavage furrow. **B.** Representative images of mitotic cell surface and cell-cell contact localization of PCDH7 in FRAP analysis. Relative timing after photo-bleaching is shown in minutes. Yellow arrows indicate the photo-bleached region. **C.** Western blotting analysis of PCDH7 protein levels in CRISPR based knockout cells. Single colonies of different guide RNAs (sg1, sg5, sg6) against PCDH7 are analyzed. Non-treated HeLa S3 (Hela parental) and non-targeting sgRNA (sgNT) treated cells are used as control. Actin was used as the loading control. **D.** Western blotting analysis of PCDH7 protein levels in rescue cells. Control guide RNA (sgNT) and PCDH7 guide RNAs (PCDH7 sg1, sg5) treated cells were analyzed in parallel with rescue cells after viral transduction of PCDH7:eGFP (sgNT+PCDH7:GFP, PCDH7 sg1+PCDH7:GFP, PCDH7 sg5+PCDH7:GFP) and control (PCDH7 sg1+emptyGFP, PCDH7 sg5+emptyGFP) constructs. The upper band corresponds to fusion protein (PCDH7:eGFP, 116+27) while the lower band corresponds to PCDH7 (116). Actin was used as the loading control.

**Figure S2.**
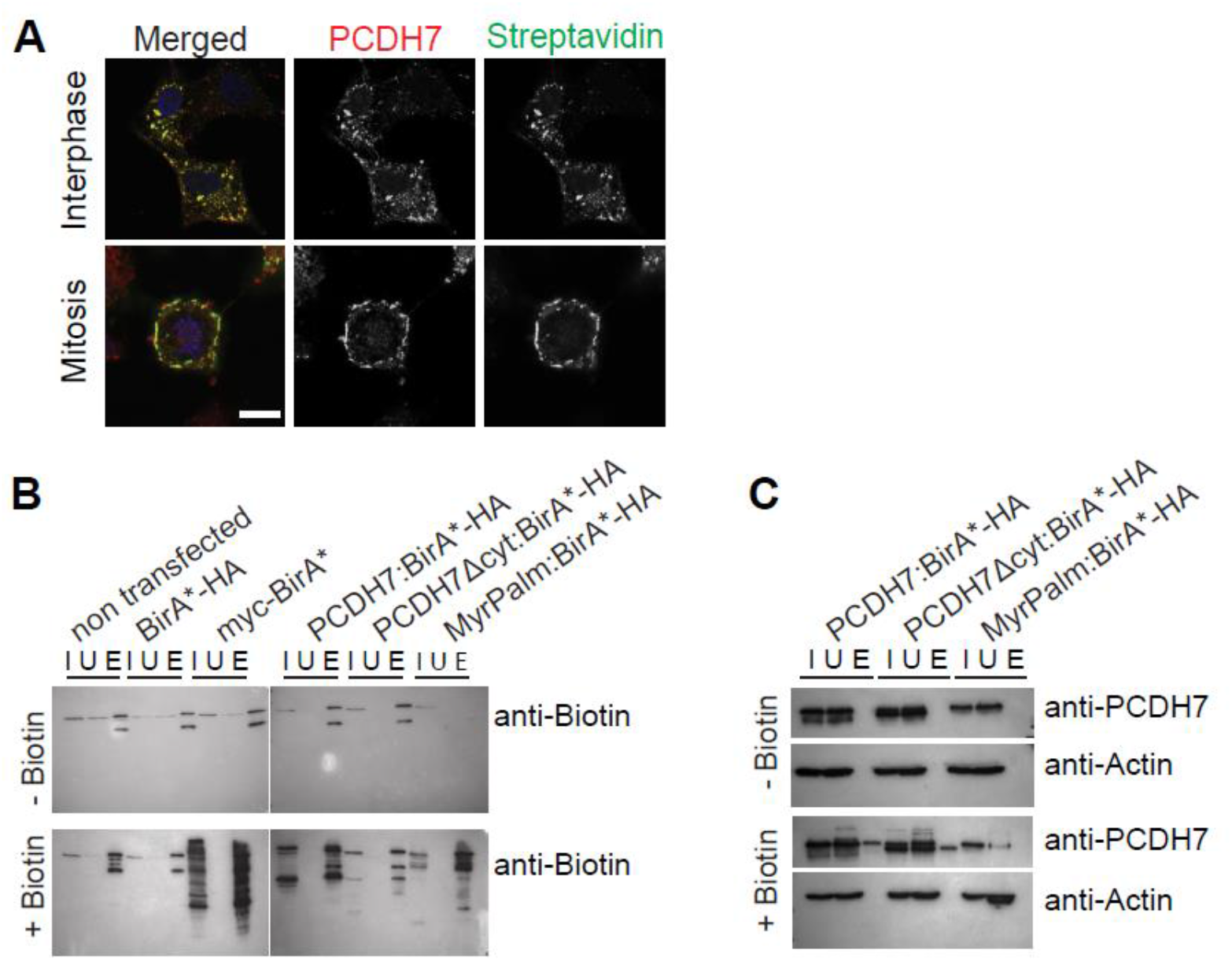
PCDH7:BirA* expression exhibits efficient and specific biotinylation. **A.** The cell cycle-specific localization of PCDH7:BirA*-HA (PCDH7 BioID construct) in interphase (top) and mitosis (bottom) cells, fixed and stained against PCDH7 antibody (red) and fluorescence conjugated streptavidin (green), DAPI in blue. **B.** Western blotting analysis of BioID constructs expressing cells’ biotinylation affinity using anti-biotin antibodies. Non-transfected, empty vector-transfected (BirA*-HA and mycBirA*) cells are used as a control (left). PCDH7:BirA*-HA (PCDH7 BioID), PCDH7Δcyt:BirA*-HA (truncated PCDH7 BioID), myrPalm:BirA*-HA (MyrPalm BioID, an unrelated construct) are analyzed (right). Upper blot; Cells not supplied with biotin. Lower blot; Cells supplied with biotin. In the absence of biotin, only a basal level of biotinylation was observed for all cell lines. The inputs (I) obtained from PCDH7 BioID, truncated PCDH7 BioID and MyrPalm BioID transfected cells demonstrated different biotinylation patterns as expected. The biotinylated proteins were successfully enriched at the elute fractions (E) of each sample after streptavidin affinity pulldown. **C.** Western blotting analysis of BioID constructs expressing cells against anti-PCDH7 and anti-actin antibodies. Both PCDH7 and truncated PCDH7 (PCDH7Δcyt) are enriched in the elute fractions (E) after streptavidin pulldown only in the biotin-supplied cells (bottom). No PCDH7 enrichment is observed in the cells not supplemented with biotin (top). The full-length PCDH7 was only present in the elute (E) fraction of PCDH7 BioID transfected cells. The absence of actin in the elute fractions suggested the lack of cytosolic contamination (bottom panel). Scale bar: 10 µm

**Figure S3.**
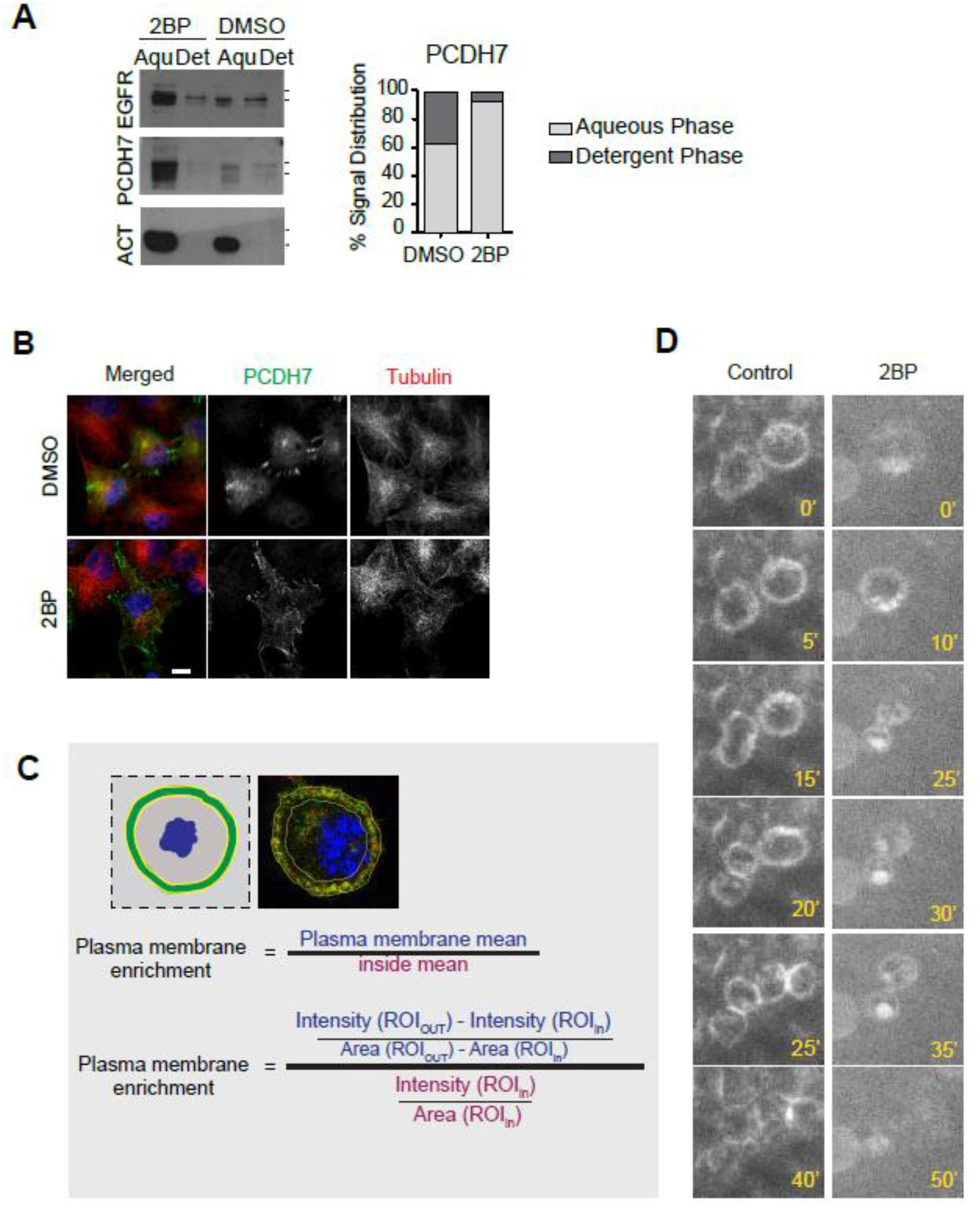
Characterization of 2BP inhibitor treatment. **A. 2BP changes the phase of PCDH7 in Triton X-114 extraction**. Western blotting analysis of the effect of 2BP on the hydrophobicity of PCDH7. Aqueous (Aqu) and detergent (det) phases are separated using Triton X-114 extraction and blotted against PCDH7, EGFR (a palmitoylated plasma membrane protein), and Actin (a cytoplasmic protein) (left). 2BP treatment increased the amount of the EGFR, a known palmitoylated protein in the aqueous phase when compared with the DMSO control. A similar effect was also observed for the PCDH7. The quantification of the PCDH7 distribution among aqueous and detergent phases in control (DMSO) and 2BP treated cells (right). **B.** PCDH7 (green, anti-GFP) localization in control (DMSO) (top) and palmitoylation inhibitor, 2BP treated (bottom) interphase cells those are stably expressing PCDH7-GFP-BAC. Microtubules (anti-α-Tubulin) are shown in red and DNA (DAPI) in blue (left). **C.** Calculation of plasma membrane enrichment. Illustration (left) and a representative fluorescence image (right) for the analyzed region of interest (ROIs) where the plasma membrane intensities are measured and normalized to inside. Corresponding formula is presented below. **D.** Live imaging snapshots of the dividing PCDH7-GFP-BAC expressing cells in the control (top) and 2BP treated (bottom) cells. Relative timing after mitotic rounding is shown in minutes. Scale bar: 10 µm

**Figure S4.**
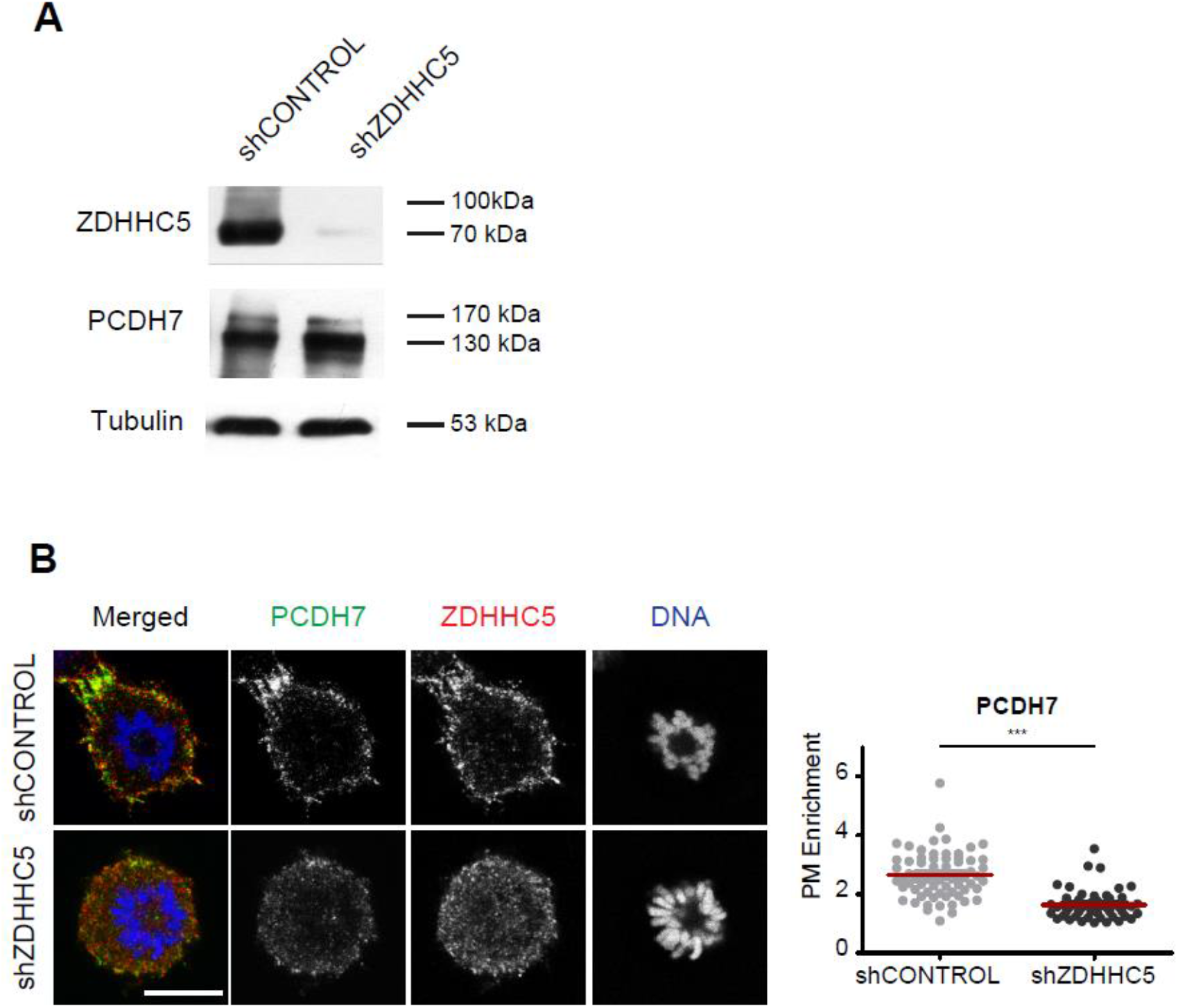
shRNA knockdown of ZDHHC5 perturbs the mitotic cell surface localization of PCDH7. **A.** Western blotting analysis of shZDHHC5 and shCONTROL cells. shZDHHC5 treatment successfully depleted ZDHHC5 while PCDH7 levels remain unchanged. Tubulin was used as the loading control. **B**. PCDH7 (green, anti-GFP) localization in shCONTROL and shZDHHC5 treated mitotic cells that are stably expressing PCDH7-GFP-BAC. Cells are synchronized to monopolar mitosis. ZDHHC5 is shown in red (anti-ZDHHC5) and DNA in blue (DAPI) (left). Quantification of the plasma membrane enrichment of PCDH7 during mitosis in shCONTROL (n=68) and shZDHHC5 (n=52) treated cells (right). Statistics used *unpaired two-tailed t-test*. Scale bar: 10 µm; ***: p<0.001

**Figure S5.**
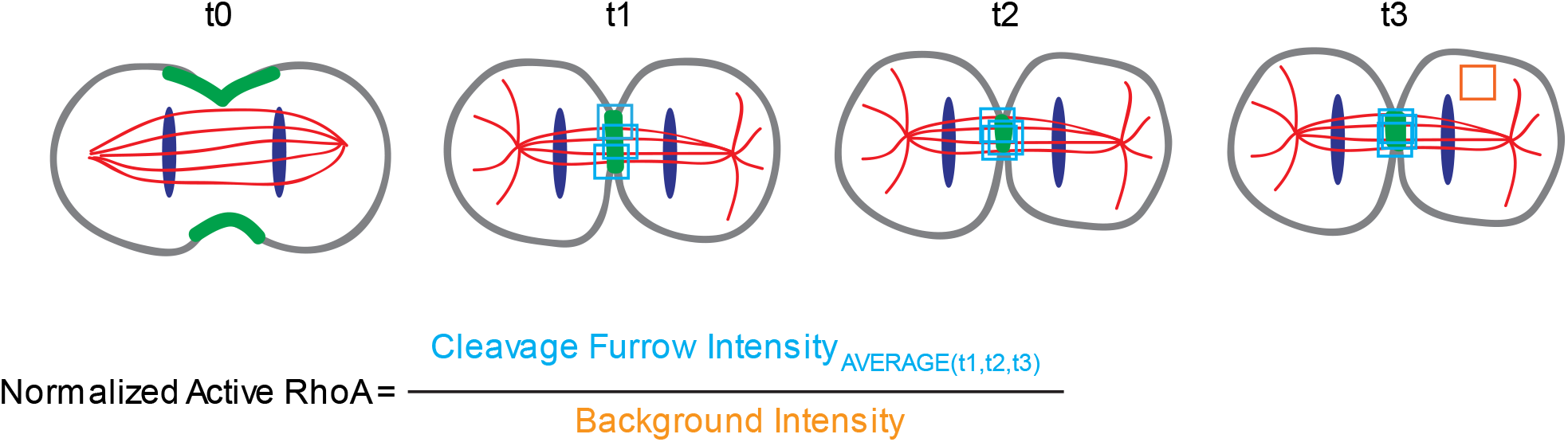
Illustration of the quantification of active RhoA levels at the cleavage furrow. A fixed-sized region of interest (ROIs) was used for all measurements and active RhoA intensity was calculated by taking the average of three random ROIs that cover the RhoA signal at the cleavage furrow. For Normalized Active-RhoA (Figure 8B); average RhoA intensity was divided by the cytoplasmic signal.

